# Progranulin haploinsufficiency remodels the cerebral microvasculature and neurovascular unit

**DOI:** 10.64898/2026.07.20.738719

**Authors:** Supriya Chakraborty, Madeleine Weick, Sofia Andrea Franciosa, Zeynab Tabrizi, Charlotte Swinehart, Gabriela Rodriguez Moore, Dylan Lee, Nairuti Nikhil Bhatt, Alejandra Cano, Poorva Poorva, Silvia Torices, Ruslan Rust, Oliver Bracko

## Abstract

Progranulin (PGRN) deficiency is a major genetic cause of frontotemporal dementia (FTD), yet its impact on cerebrovascular function remains understudied. Here, we show that PGRN deficiency contributes to cerebral microvascular perfusion and induces alterations within the neurovascular unit. *In vivo* two-photon imaging revealed increased capillary stalling and in cerebral blood flow (CBF), driven in part by increased leucocyte-capillary interactions and elevated endothelial ICAM-1 expression. Transcriptomic profiling of isolated cerebral microvessels demonstrated coordinated upregulation of immune and extracellular matrix pathways alongside suppression of angiogenic and stress-response programs, indicative of endothelial activation. Cross-species analyses identified partial conservation of these vascular signatures in endothelial cells from human FTD-GRN patients, associated with dysregulated angiogenic and inflammatory signaling. Despite altered tight junction organization and reduced solute carrier transporter expression, blood–brain barrier (BBB) permeability remained largely intact, suggesting functional rather than structural BBB disruption, as well as. These vascular changes were accompanied by broad alterations in the morphology of astrocytes, pericytes, and microglial cells. Here we determined a novel role for progranulin in cerebrovascular homeostasis and established microvascular dysfunction as a key driver of FTD-GRN pathophysiology.

## INTRODUCTION

Frontotemporal dementia (FTD) is the second most common cause of dementia in adults under the age of 65, affecting an estimated 60,000–90,000 individuals in the United States but is producing a disproportionate burden of disability in midlife ^1,2^. FTD is clinically heterogeneous, presenting most often with progressive behavioral disinhibition, executive dysfunction, or language impairment ^3^. Approximately 30% of cases are inherited, with the majority of familial disease attributable to germline mutations in three genes: progranulin (*GRN*), the hexanucleotide repeat expansion in *C9orf72*, and microtubule-associated protein tau (*MAPT*) ^4–8^. *GRN*-associated FTD (FTD-*GRN*) is driven by haploinsufficiency: a single loss-of-function allele reduces circulating and tissue progranulin (PGRN) to roughly half of normal levels and is sufficient to cause disease, making progranulin biology a focal point of mechanistic and therapeutic investigation ^5^.

In the brain, PGRN is expressed predominantly by neurons and microglia ^9^, where it is trafficked to lysosomes and is secreted into the extracellular space ^10,11^. Its established functions include support of lysosomal homeostasis, neuroprotection, and regulation of neuronal growth and maturation ^11,12^. Beyond these neuronal and microglial roles, PGRN has been implicated in modulating inflammatory signaling, extracellular matrix remodeling, and tissue-repair processes that influence vascular and endothelial function in peripheral tissues ^13,14^. Specifically, PGRN attenuates pro-inflammatory cytokine signaling and modulates leukocyte–endothelial interactions, raising the possibility that P*GRN* deficiency contributes to FTD-*GRN* pathology through both cerebrovascular and neuronal mechanisms ^13,14^. Because PGRN haploinsufficiency is the proximate cause of FTD-*GRN*, progranulin-deficient mice, heterozygous and homozygous, are widely used to dissect these mechanisms *in vivo* ^15,16^.

Despite these circumstantial links between PGRN and vascular biology, mechanistic studies in FTD have overwhelmingly focused on neurons, microglia, and lysosomal pathways, leaving the cerebrovascular compartment comparatively understudied ^17^. A single-nucleus transcriptomic analysis of postmortem FTD-*GRN* cortex identified structural and molecular alterations in endothelial cells, pericytes, and the blood–brain barrier (BBB), indicating direct vascular involvement in disease progression ^18^. In parallel, two recent studies in murine and cell culture systems reported impaired BBB function in TDP-43 proteinopathy and *C9orf72 E*xpansion models ^19,20^, establishing cerebrovascular dysfunction as an emerging feature across the FTD spectrum. Whether analogous changes occur in FTD-*GRN* and whether PGRN haploinsufficiency itself is sufficient to engage these mechanisms *in vivo have* not been directly tested ^21,22^.

The BBB is essential for separating circulating blood components from the brain parenchyma ^23–25^. It is formed by specialized brain endothelial cells joined by continuous tight junctions, with supporting contributions from pericytes, astrocytic endfeet, and the basement membrane, and it tightly regulates the passage of solutes, immune cells, and circulating neurotoxic factors into the central nervous system ^24,25^. The integral membrane proteins Claudin-5 and Occludin are central to tight-junction integrity ^26–28^. Claudin-5 is a 23 kDa protein expressed at brain endothelial cell junctions, where it primarily forms homophilic interactions with Claudin-5 on adjacent cells and heterophilic interactions with other claudin family members, such as Claudin-1 and Claudin-3 ^28–30^. Occludin, a 65 kDa integral membrane protein, associates laterally with claudins and binds the cytoplasmic scaffold ZO-1 to stabilize tight-junction architecture and restrict paracellular permeability ^31^. Loss of BBB integrity is a recurrent feature of neurodegenerative diseases, including Alzheimer’s Disease, Amyotrophic Lateral Sclerosis, Huntington’s Disease, and multiple sclerosis, where it is associated with neuroinflammation, edema, and cognitive decline, positioning the barrier as a key checkpoint linking systemic and central pathology ^32–35^.

A study reported reduced cerebral blood flow (CBF), measured by arterial spin-labeling MRI in midlife and presymptomatic *GRN* mutation carriers ^36^, implicating cerebrovascular dysfunction at the early stage of disease. At the capillary level, we and others have previously shown that leukocytes can adhere to and stall in brain capillaries, contributing to hypoperfusion, neuroinflammation, and cognitive impairment in Alzheimer’s Disease models ^37,38^. Because capillaries are the principal site of nutrient and gas exchange, ionic balance, and clearance of metabolic waste, even modest disruptions to capillary flow can compromise neuronal health ^37–40^. These microvascular processes are coordinated within the neurovascular unit (NVU), in which endothelial cells, pericytes, astrocytes, neurons, and microglia interact at the capillary level to regulate CBF, maintain BBB integrity, and clear metabolic waste^41–44^. Whether analogous microvascular dysfunction, capillary stalling, endothelial activation, and impaired perfusion occur in FTD-*GRN*, and whether such changes are accompanied by broader structural NVU remodeling, has not been directly examined.

Here, we test the hypothesis that progranulin haploinsufficiency drives cerebrovascular dysfunction and remodeling of the NVU *in vivo*. We combine intravital two-photon imaging of cortical microvasculature, transcriptional profiling of isolated cerebral microvessels, structural and functional assessment of the BBB, NVU cellular morphometry, and protein-level interrogation of vascular signaling in heterozygous (PGRN^+/−^) and homozygous (PGRN^−/−)^ progranulin-deficient mice. Cross-species comparison with human FTD-*GRN* endothelial transcriptomes anchors these findings to disease relevance. Together, the data position the cerebral endothelium and NVU as core components of FTD-*GRN* pathology and offer a mechanistic context for the cerebrovascular signatures observed in human *GRN* mutation carriers.

## RESULTS

### Progranulin deficiency disrupts cerebral microvascular homeostasis and promotes leukocyte-mediated capillary obstruction

To determine whether progranulin deficiency alters cerebral microvascular function *in vivo*, we performed two-photon fluorescence (2PEF) imaging through chronic cranial windows in aged WT, PGRN⁺/⁻, and PGRN⁻/⁻ mice (Fig. 1A). Three-dimensional reconstruction of cortical vascular networks labeled with Texas Red dextran revealed reduced vessel density in progranulin-deficient mice compared with WT controls (Fig. 1B), consistent with microvascular rarefaction. Quantification of stalled capillary segments demonstrated a significant increase in both PGRN⁺/⁻ and PGRN⁻/⁻ mice relative to WT animals (Fig. 1C and Supplementary Fig. 1). Because capillary stalls can arise from distinct cellular mechanisms, we next characterized the cellular composition of stalled vessels. Whereas stalls in WT and PGRN⁺/⁻ mice were predominantly associated with red blood cells, PGRN⁻/⁻ mice exhibited a shift toward leukocyte-associated capillary stalls (Fig. 1D).

**Figure 1:**
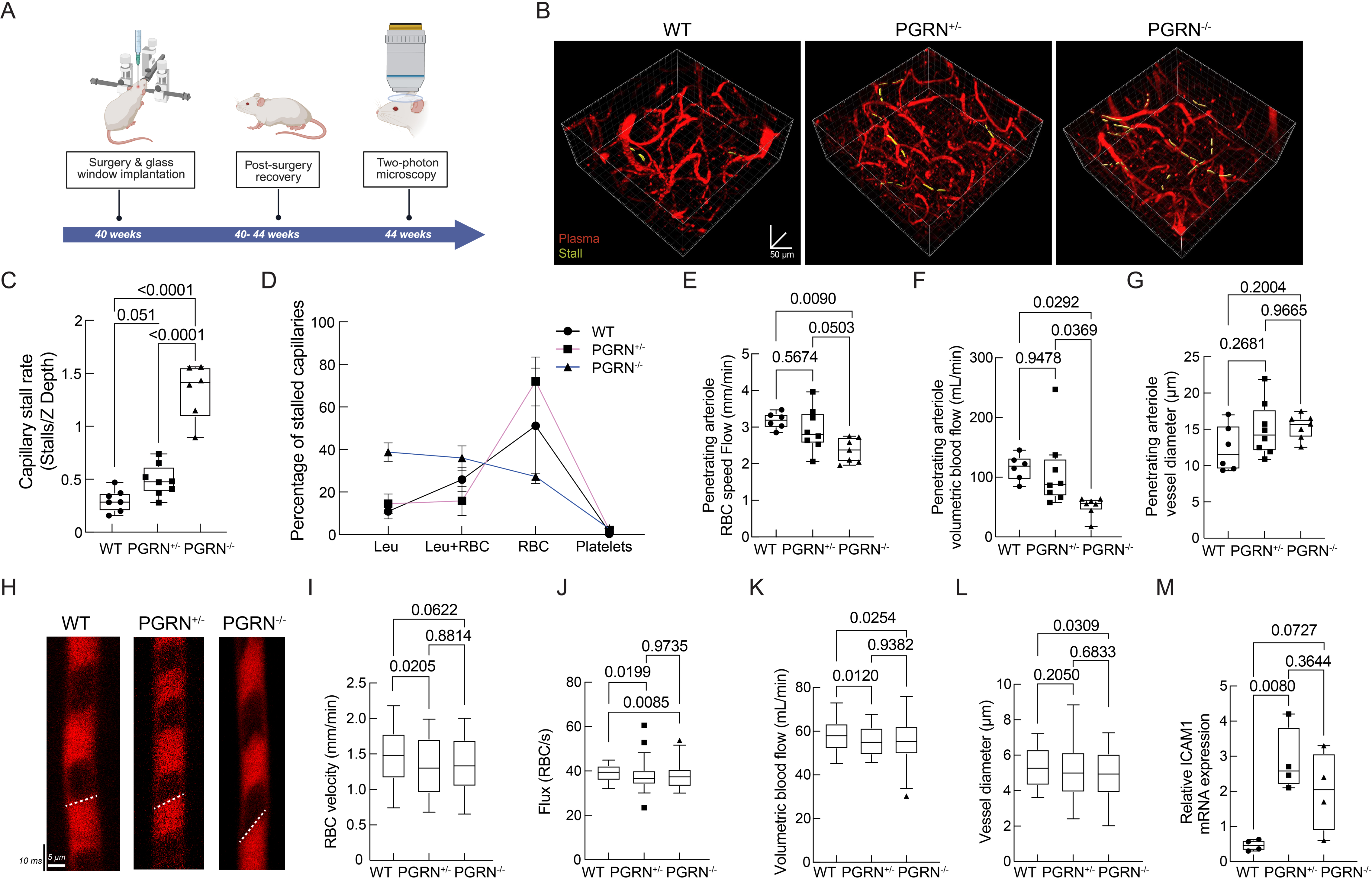
Progranulin deficiency causes microvascular rarefaction, increased capillary stalling, and impaired cerebral perfusion with endothelial activation. (A) Overview and timeline of craniotomy surgery and multiphoton imaging. (B) Three-dimensional reconstructions of *in vivo* 2PEF image stacks from cortical vasculature labeled with 70 kDa Texas Red dextran (red, plasma); stalled capillaries are highlighted in yellow. (C) Quantification of capillary stall rate across genotypes WT, PGRN⁺/⁻, and PGRN⁻/⁻. (D) Cellular composition of capillary stalls by genotype; stalls in WT mice are predominantly RBC-associated, whereas PGRN⁻/⁻ mice exhibit a proportional shift toward leukocyte-associated stalls, indicative of increased leukocyte–endothelial interaction. Penetrating arteriole RBC velocity (E), volumetric blood flow (F), and vessel diameter (G) across genotypes. Penetrating arterioles analyzed: WT (n = 27), PGRN⁺/⁻ (n = 31), and PGRN⁻/⁻ (n = 19). (H) Representative line scans from WT, PGRN⁺/⁻, and PGRN⁻/⁻ showing capillary blood flow analysis. (I–L) Capillary RBC velocity (I), RBC flux (J), volumetric blood flow (K), and vessel diameter (L). Capillaries analyzed: WT (n = 106), PGRN⁺/⁻ (n = 113), and PGRN⁻/⁻ (n = 125). Groups and sex: PGRN⁻/⁻ (n = 7; 4M/3F), PGRN⁺/⁻ (n = 7; 3M/4F), WT (n = 6; 4M/2F). Statistical comparisons were performed using Kruskal–Wallis one-way ANOVA with Dunn’s post hoc test.; (M) ICAM-1 mRNA expression in isolated cerebral microvessels, normalized to GAPDH; expression is elevated in progranulin-deficient mice. Data are presented as individual values with group summary statistics unless otherwise indicated.

Given the increased prevalence of capillary stalling, we next assessed whether this was accompanied by impaired cerebral perfusion. Because carriers of pathogenic *GRN* mutations exhibit cerebral hypoperfusion ^36^, we quantified cerebral blood flow (CBF) in penetrating arterioles and cortical capillaries using 2PEF line-scan imaging. PGRN-deficient mice displayed progressive reductions in penetrating arteriole RBC velocity (Fig. 1E) and volumetric blood flow (Fig. 1F), whereas vessel diameter remained unchanged across genotypes (Fig. 1G). Similar reductions were observed within the capillary network, where progranulin-deficient mice exhibited decreased RBC velocity (Fig. 1H, I), RBC flux (Fig. 1J), and volumetric blood flow (Fig. 1K). In contrast to penetrating arterioles, capillary diameter was significantly reduced in PGRN⁻/⁻ mice relative to WT controls (Fig. 1L). To investigate a potential mechanism underlying the increased leukocyte-associated stalling phenotype, we measured mRNA expression of the endothelial adhesion molecule ICAM-1 in isolated cerebral microvessels. ICAM-1 expression was significantly elevated in progranulin-deficient mice (Fig. 1M), consistent with enhanced endothelial activation and increased leukocyte–endothelial interactions. Collectively, these findings demonstrate that progranulin deficiency is associated with reduced microvascular density, increased capillary stalling, impaired CBF, and elevated endothelial activation. The shift toward leukocyte-associated capillary stalls together with increased ICAM-1 expression suggests that inflammatory vascular mechanisms contribute to cerebral hypoperfusion in progranulin-deficient mice.

### Microvessel transcriptomics reveals coordinated immune activation and extracellular matrix remodeling in progranulin deficiency

To define molecular mechanisms underlying cerebrovascular dysfunction in progranulin deficiency, we performed RNA sequencing of isolated cerebral microvessels from WT, PGRN^+/−^, and PGRN^−/−^ mice (Fig. 2A). Comparative analysis demonstrated substantial overlap in expressed genes across genotypes, while differential expression analysis identified 180 DEGs between WT and PGRN^+/−^ microvessels, 316 DEGs between WT and PGRN^−/−^ microvessels, and 32 DEGs between PGRN^+/−^ and PGRN^−/−^ microvessels (Fig. 2B and Supplementary Table 1). Volcano plot analysis revealed widespread transcriptional alterations in PGRN^+/−^ microvessels relative to WT controls, including both upregulated and downregulated genes (Fig. 2C). To determine the biological processes associated with these changes, we performed vascular-focused Gene Ontology enrichment analysis. Upregulated pathways were predominantly associated with extracellular matrix organization, extracellular structure organization, tissue remodeling, and extracellular matrix assembly (Fig. 2D). In contrast, pathways related to angiogenesis, vascular development, oxidative stress responses, reactive oxygen species metabolism, and tumor necrosis factor production were significantly downregulated, indicating broad alterations in vascular homeostatic signaling. To further resolve the molecular networks underlying these transcriptional changes, we performed protein–protein interaction analysis using DEGs from progranulin-deficient microvessels. This analysis identified several interconnected functional modules enriched for immune activation and lymphocyte signaling, oxidative stress and NADPH oxidase pathways, antigen presentation and MHC signaling, vascular remodeling and adhesion, chemokine-mediated leukocyte recruitment, and extracellular matrix/collagen organization (Fig. 2E–J). Collectively, these findings demonstrate that progranulin deficiency induces transcriptional reprogramming within the cerebral microvasculature, characterized by coordinated activation of immune-associated pathways and dysregulation of extracellular matrix remodeling programs.

**Figure 2.**
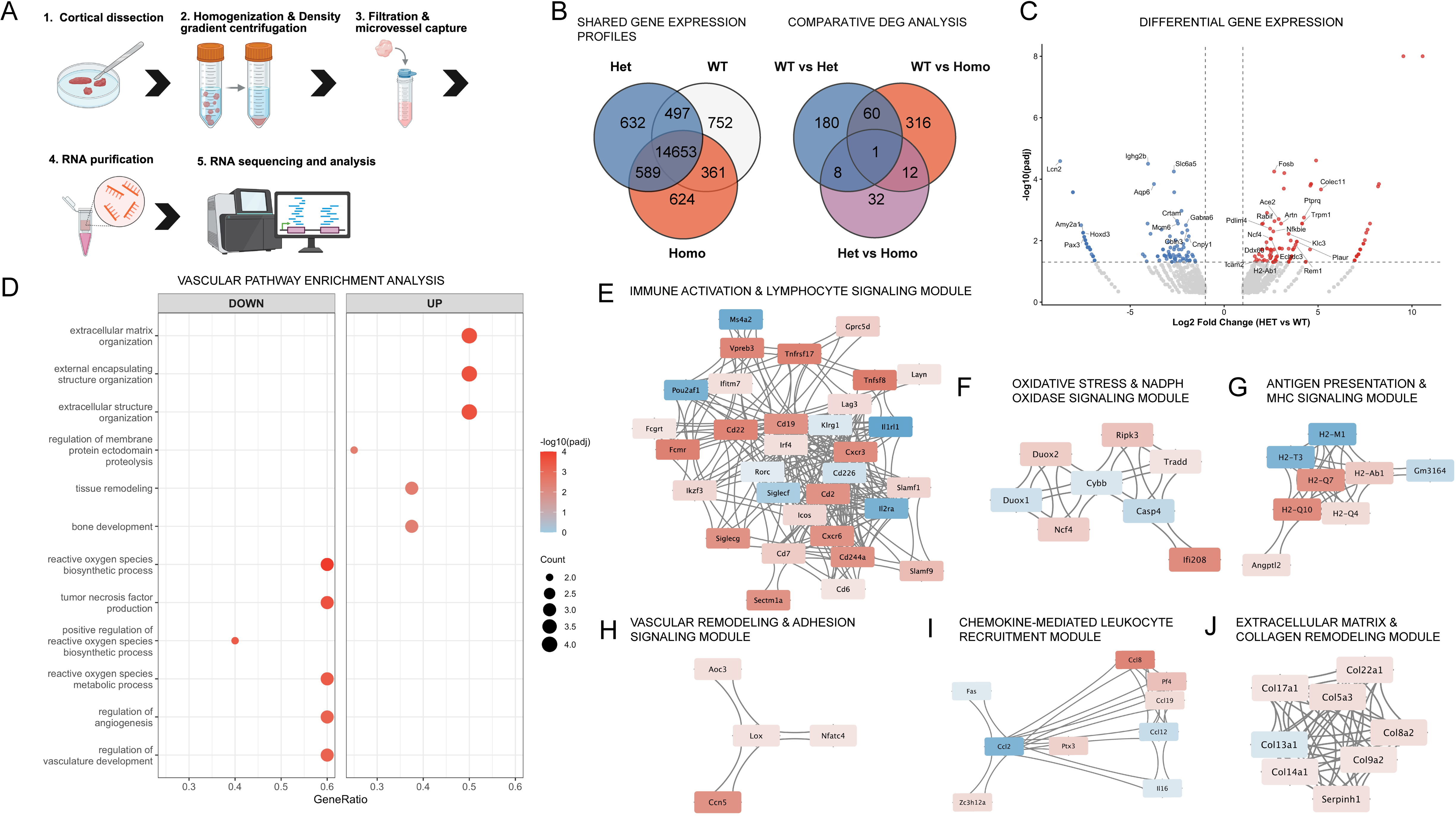
RNA sequencing of isolated cerebral microvessels reveals immune activation and extracellular matrix remodeling in progranulin-deficient mice. (A) Workflow of microvessel isolation. (B) Venn diagrams depicting overlap of expressed genes (left) and significantly differentially expressed genes (DEGs; adjusted p < 0.05, right) across WT, PGRN^+/−^, and PGRN^−/−^ cerebral microvessel datasets. (C) Volcano plot of DEGs in PGRN^+/−^ versus WT microvessels. Upregulated genes are shown in red, downregulated in blue, and non-significant genes in gray. Xist was excluded from visualization. (D) Vascular-focused Gene Ontology analysis of DEGs in PGRN^+/−^ versus WT microvessels. (E–J) Protein–protein interaction network modules: immune activation and lymphocyte signaling (E), oxidative stress and NADPH oxidase pathways (F), antigen presentation and MHC signaling (G), vascular remodeling and adhesion signaling (H), chemokine-mediated leukocyte recruitment (I), extracellular matrix and collagen organization (J).

### Cross-species endothelial transcriptomic analysis reveals conserved vascular remodeling programs in progranulin deficiency

To determine whether endothelial transcriptional alterations observed in progranulin-deficient mice recapitulate those present in human FTD-GRN, we compared differential expression signatures derived from human capillary endothelial pseudobulk transcriptomes ^18^ with endothelial differential expression profiles obtained from PGRN^+/−^ and PGRN^−/−^ mouse cerebral microvessels. A heatmap of the capillary endothelial genes exhibiting the strongest transcriptional separation between FTD-GRN and control donors demonstrated clear segregation of disease and control samples, indicating robust disease-associated transcriptional remodeling of the endothelial compartment (Fig. 3A). We next assessed cross-species directional concordance of orthologous gene expression changes between human and mouse datasets. Comparison of human FTD-GRN capillary endothelial signatures with PGRN^+/−^ mice identified 107 shared orthologous genes, of which 52 exhibited concordant regulation and 55 showed opposite-direction changes (Fig. 3B). Similarly, comparison with PGRN^−/−^ mice identified 137 shared orthologous genes, including 64 concordantly regulated genes and 73 oppositely regulated genes (Fig. 3C). These findings demonstrate partial conservation of progranulin-dependent endothelial transcriptional responses across species, with both mouse models recapitulating distinct components of the human endothelial disease signature.

**Figure 3.**
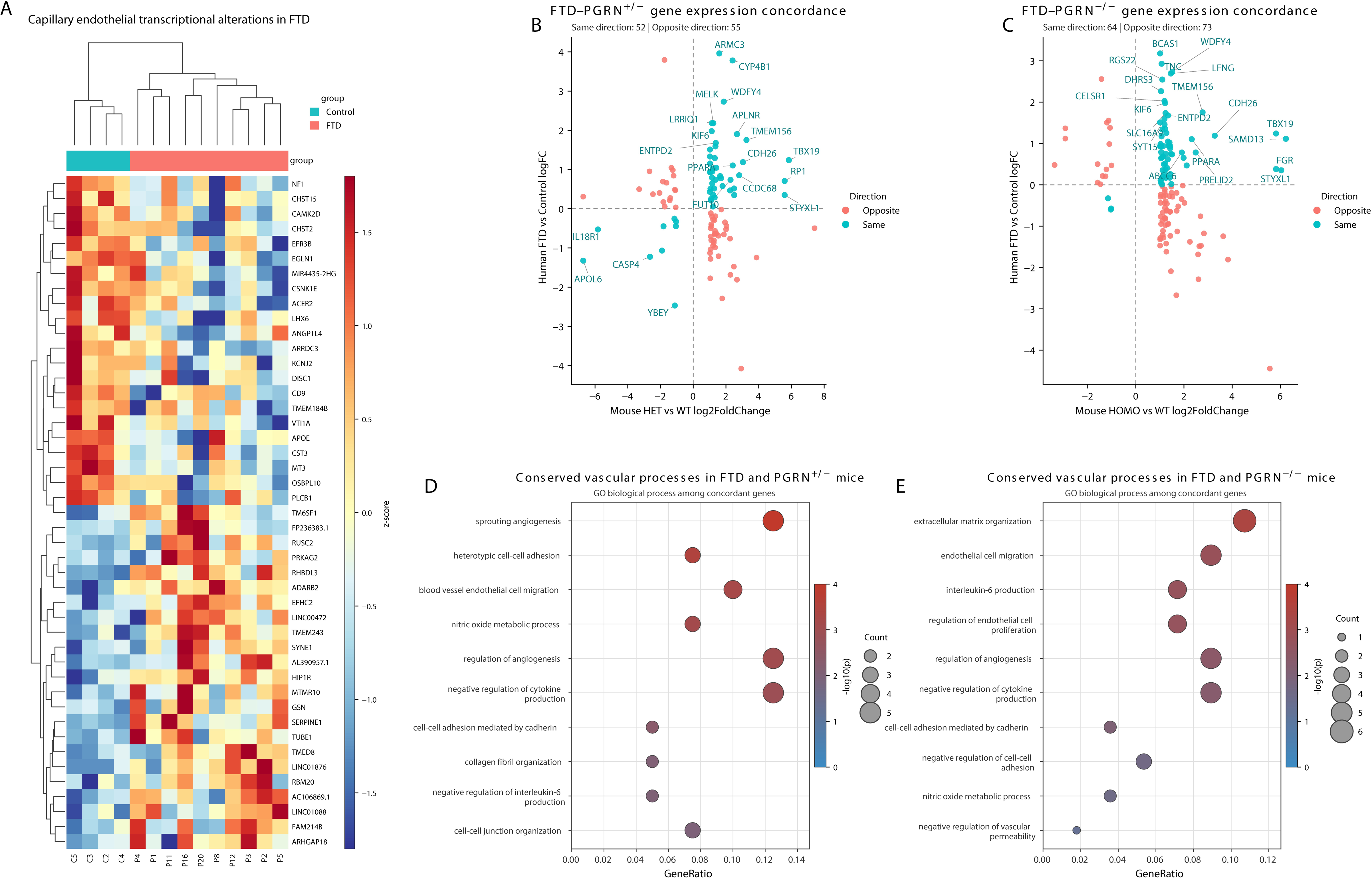
Cross-species comparison of endothelial transcriptional responses to progranulin deficiency. (A) Heatmap of the 45 capillary endothelial genes showing the strongest transcriptional separation between human FTD-GRN and control donors. (B–C) Scatterplots comparing log₂ fold changes of shared orthologous genes between human FTD-GRN and mouse PGRN^+/−^ (B) or PGRN^−/−^ (C) microvessel RNA-seq datasets. Points are colored according to directional concordance, and dashed lines indicate zero log₂ fold change on each axis. Gene Ontology Biological Process enrichment analysis of cross-species concordant genes highlights representative conserved vascular biological processes in PGRN^+/−^ (D) and PGRN^−/−^ (E) mice. Dot size represents gene count, and color indicates −log₁₀(p-value).

To investigate the biological processes shared between species, Gene Ontology biological process enrichment analysis was performed using concordantly regulated genes identified in both human FTD-GRN and mouse datasets. Concordant genes shared between human FTD-GRN and PGRN^+/−^ mice were enriched for vascular remodeling processes, including sprouting angiogenesis, endothelial cell migration, regulation of angiogenesis, nitric oxide metabolic processes, collagen fibril organization, and cell-cell junction organization (Fig. 3D). Similarly, concordant genes shared between human FTD-GRN and PGRN^−/−^ mice were enriched for extracellular matrix organization, endothelial cell migration, regulation of endothelial cell proliferation and angiogenesis, vascular permeability, nitric oxide metabolic processes, and inflammatory signaling, including regulation of interleukin-6 production (Fig. 3E). These results demonstrate that progranulin deficiency induces a conserved endothelial remodeling program across mouse and human, characterized by alterations in angiogenesis, extracellular matrix organization, endothelial migration, vascular homeostasis, and inflammatory signaling, supporting the translational relevance of the vascular mechanisms identified in the PGRN mouse model.

### Progranulin deficiency disrupts vascular maintenance signaling and reduces cortical vessel density

Microvessel transcriptomic analysis identified downregulation of pathways associated with angiogenesis and vascular development in PGRN^+/−^ mice, while cross-species comparison showed that angiogenesis, endothelial migration, extracellular matrix organization, cell adhesion pathways, and related vascular processes were shared components of the endothelial response in mouse models and human FTD-GRN (Figs. 2, 3). We therefore investigated whether these transcriptional alterations were accompanied by changes in angiogenesis-associated proteins and cortical vascular architecture. Angiogenesis-focused protein profiling of cortical lysates revealed broad alterations in vascular-associated proteins in PGRN^+/−^ mice relative to WT controls (Fig. 4A, B). Among the most significantly altered analytes, Osteopontin and NOV/CCN3 were reduced in PGRN^+/−^ cortical lysates (Fig. 4C, D and Supplementary Fig. 2A). Ranking of array-derived proteins and volcano plot analysis further demonstrated that several angiogenesis-associated factors were differentially expressed, with Osteopontin and NOV/CCN3 among the strongest reductions detected in the heterozygous mice (Fig. 4E, F and Supplementary Fig. 2B, C). To assess a major vascular trophic mediator independently, we quantified VEGF-A in cortical lysates by ELISA. VEGF-A concentrations were significantly reduced in PGRN^+/−^ mice relative to WT controls (Fig. 4G), providing additional protein-level evidence of altered vascular maintenance signaling. We next examined whether these molecular changes were accompanied by alterations in the cortical vascular network. CD-31 immunostaining revealed a reduced number of cortical blood vessels in PGRN^+/−^ mice than in WT controls (Fig. 4H, I). Because microvessel ICAM-1 transcript expression was elevated and leukocyte-associated capillary stalls were increased in progranulin-deficient mice, we also measured soluble ICAM-1 and VCAM-1 in cortical lysates. Neither ICAM-1 nor VCAM-1 protein concentrations differed significantly between WT and PGRN^+/−^ mice, but both showed a trend towards increased levels in PGRN^+/−^ mice (Fig. 4J, K). This series of experiments extends the transcriptomic findings by demonstrating reduced levels of several vascular maintenance-associated proteins together with decreased cortical vessel density in PGRN^+/−^ mice. Thus, progranulin haploinsufficiency is associated with coordinated molecular and structural remodeling of the cerebral vasculature.

**Figure 4.**
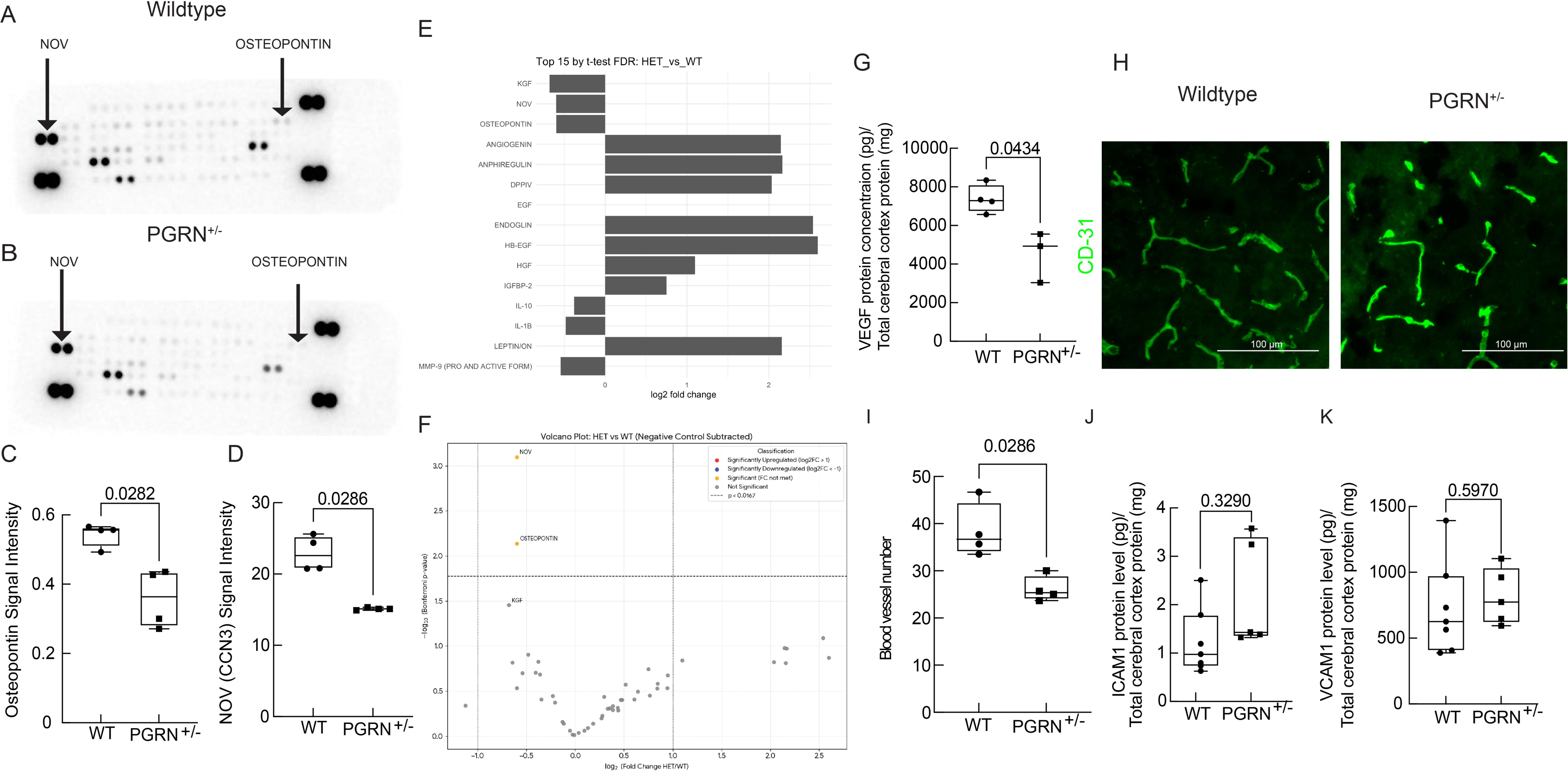
Progranulin haploinsufficiency alters vascular maintenance-associated proteins and reduces cortical vessel density. Representative angiogenesis antibody array membranes generated from cortical lysates of WT (A) and PGRN^+/−^ (B) mice. Arrows indicate the duplicate spots corresponding to NOV/CCN3 and Osteopontin. (C–D) Quantification of normalized array signal intensity for Osteopontin (C) and NOV/CCN3 (D), demonstrating reduced abundance in PGRN^+/−^ cortical lysates relative to WT controls. (E) Ranking of the 15 angiogenesis-associated proteins showing the highest differences between PGRN^+/−^ and WT samples. Values are presented as log₂ fold change. (F) Volcano plot of differential protein abundance in PGRN^+/−^ versus WT cortical lysates following background subtraction and normalization. Dashed lines indicate the statistical and fold-change thresholds used for visualization. (G) ELISA quantification of VEGF-A in cortical lysates from WT and PGRN^+/−^ mice, normalized to total protein content. (H) Representative images of lectin-labeled cortical blood vessels in WT and PGRN^+/−^ mice. Scale bar, 100 μm. (I) Quantification of cortical blood vessel profiles in WT and PGRN^+/−^ mice. ELISA quantification of ICAM-1 (J) and VCAM-1 (K) in cortical lysates from WT and PGRN^+/−^ mice, normalized to total protein content. Data are presented as individual biological replicates with group summary statistics. Exact p values are indicated above the corresponding comparisons. Statistical analyses were performed as described in Methods.

### Blood–brain barrier remodeling occurs in progranulin deficiency without overt permeability defects

We next examined whether progranulin deficiency affected the structural and functional properties of the blood–brain barrier (BBB). Immunostaining of cortical microvessels revealed a significant reduction in Claudin-5 coverage along CD31-positive vessels in PGRN^+/−^ mice relative to WT controls (Fig. 5A, B), indicating altered organization of endothelial tight junctions. Occludin association with lectin-labeled vessels exhibited a similar downward trend, although the difference did not reach statistical significance (Fig. 5C, D). Consistent with these histological findings, immunoblot analysis showed reductions in total Occludin and Claudin-5 protein abundance in progranulin-deficient brain lysates, although neither change reached statistical significance (Fig. 5E, F). Together, these observations indicate remodeling of tight-junction organization rather than uniform or complete loss of junctional proteins. We next determined whether these structural changes were accompanied by increased BBB permeability. Immunostaining for endogenous IgG revealed no significant increase in parenchymal IgG immunoreactivity in PGRN^+/−^ mice relative to WT controls (Fig. 5G, H), indicating preservation of large-molecule barrier restriction under basal conditions. Similarly, sodium fluorescein extravasation did not differ significantly among WT, PGRN^+/−^, and PGRN^−/−^mice (Fig. 5I). Thus, despite altered tight-junction organization, progranulin deficiency did not produce detectable leakage to either endogenous IgG or sodium fluorescein in the assays used here. Because the BBB functions not only as a paracellular barrier but also as a specialized transport interface ^25^, we next examined endothelial transport-related transcriptional changes. Reactome enrichment analysis of differentially expressed genes from isolated cerebral microvessels identified significant alterations in solute carrier-mediated transport pathways (Fig. 5J). Heatmap analysis further demonstrated broad dysregulation of SLC transporter genes across PGRN^+/−^ and PGRN^−/−^ microvessels relative to WT samples (Fig. 5K), suggesting that endothelial transport functions may be altered even in the absence of overt tracer leakage. To further define molecular programs associated with BBB remodeling, we performed protein–protein interaction analysis of differentially expressed genes from isolated cerebral microvessels. This analysis identified three interconnected modules related to extracellular matrix remodeling and barrier organization, innate immune signaling, and coagulation or blood-derived signaling (Fig. 5L–N). The extracellular matrix remodeling and barrier disruption module showed increased Mmp14 and Timp1 expression, along with reduced Mmp12 and Pappa expression (Fig. 5L) ^45^. This mixed matrix-remodeling signature indicates altered regulation of extracellular matrix turnover rather than uniform activation or suppression of proteolysis ^46,47^. The innate immune activation module included increased Ager and Pnpla3 expression together with reduced Cd36 and Emp1 expression (Fig. 5M). These changes indicate dysregulation of inflammatory, scavenger-receptor, and membrane-associated signaling pathways, further supporting an altered microvascular immune state in progranulin deficiency ^48,49^. The coagulation and blood-derived signaling module was characterized by increased expression of Ambp, Pan, Serpinc1, and Tfpi2, along with reduced Klkb1 expression (Fig. 5N). This pattern indicates remodeling of anticoagulant, protease-inhibitory, and circulating protein-associated pathways within progranulin-deficient microvessels ^50^. In summary, progranulin deficiency remodels the BBB across several dimensions, including tight-junction organization, endothelial transporter expression, extracellular matrix regulation, innate immune signaling, and coagulation-associated pathways. However, these molecular and structural alterations occur without detectable increases in basal permeability to endogenous IgG or sodium fluorescein, indicating that BBB dysfunction in this model is characterized by endothelial reprogramming rather than overt barrier rupture ^51^.

**Figure 5:**
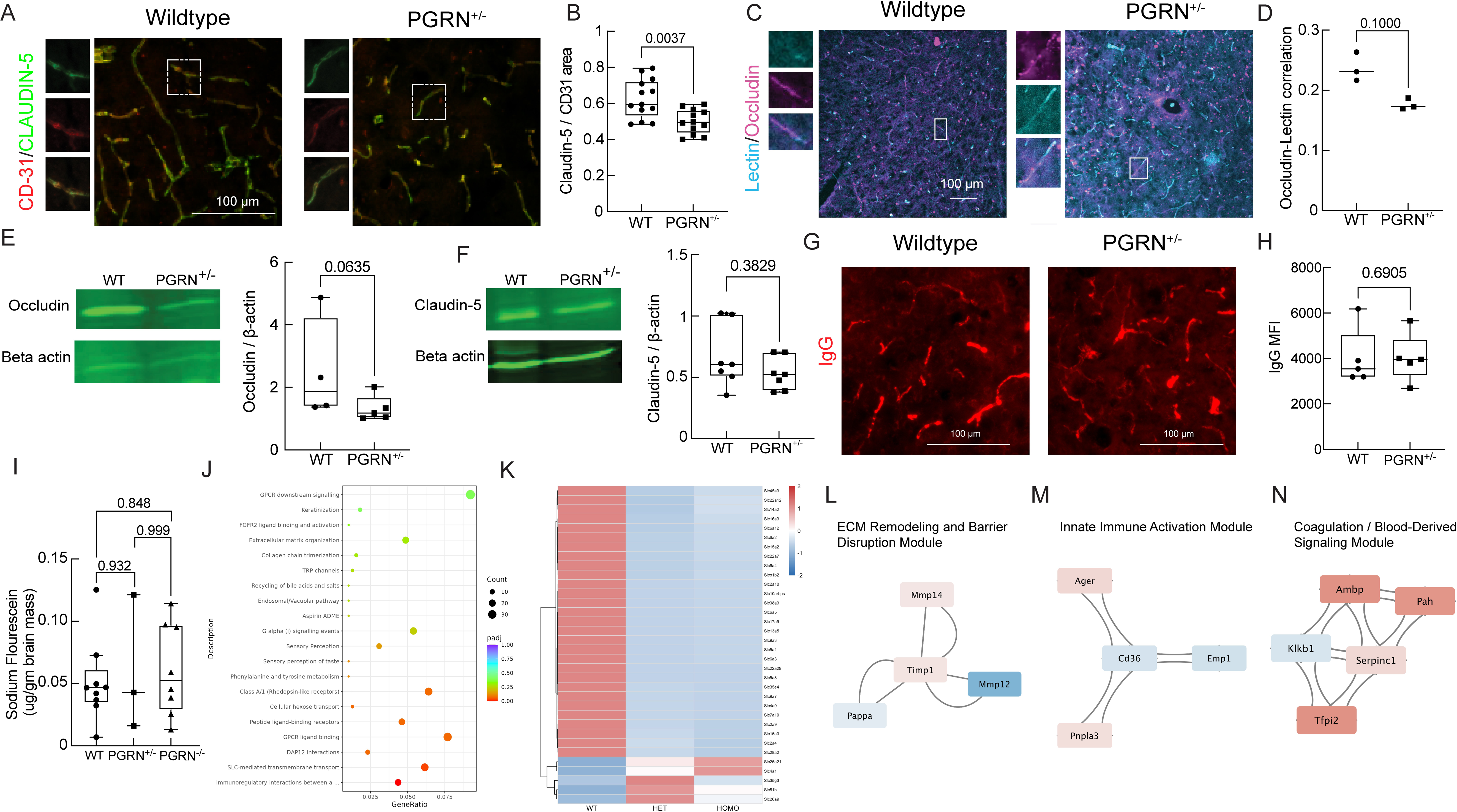
Blood–brain barrier tight junction organization and transporter expression in progranulin-deficient mice. (A) Representative immunostaining of cortical microvessels labeled with Claudin-5 (green) and CD31 (red) in WT and PGRN^+/−^ mice. Insets show magnified vessel segments. Scale bar, 100 μm. (B) Quantification of Claudin-5 coverage along CD31-positive vessels in WT and PGRN^+/−^ mice. (C) Representative immunostaining of Occludin and lectin in cortical microvessels. Insets show magnified vessel segments. (D) Quantification of Occludin association with lectin-labeled vessels in WT and PGRN^+/−^ mice. Immunoblot analysis of Occludin (E) and Claudin-5 (F) protein levels in whole-brain lysates. β-actin served as a loading control. Representative immunostaining for endogenous IgG extravasation in cortical sections as a measure of BBB permeability (G), scale bar, 100 μm, and its quantification (H). (I) Sodium fluorescein extravasation in the brain of WT, PGRN^+/−^, and PGRN^−/−^ mice. (J) Reactome pathway enrichment analysis of DEGs from cerebral microvessel RNA sequencing (PGRN^+/−^ vs WT). (K) Heatmap of SLC transporter gene expression in PGRN^+/−^, PGRN^−/−^, and WT microvessels. (L-N) Protein–protein interaction network derived from DEG analysis identifying regulatory hubs centered on TIMP1, CD36, and SERPIN family members. Data are presented as individual values with summary statistics. Statistical comparisons were performed as described in Methods.

### Neurovascular unit architecture is broadly remodeled in the progranulin-deficient cortex

To determine whether progranulin haploinsufficiency was accompanied by structural alterations in non-endothelial cellular components of the neurovascular unit (NVU). Immunolabeling of cortical astrocytes revealed marked morphological changes in PGRN^+/−^ mice. GFAP-positive astrocytes exhibited significantly reduced total branch length and fewer branches relative to WT controls (Fig. 6A–C). Sholl analysis further demonstrated decreased process intersections across radial distances from the soma, indicating reduced astrocytic arbor complexity in the progranulin-haploinsufficient cortex (Fig. 6D). Because astrocytic endfeet form a major component of the gliovascular interface and contribute to water homeostasis, BBB support, and neurovascular coupling, we next quantified AQP4-positive coverage along CD31-positive cortical vessels ^52^. Despite the reduction in overall astrocytic process complexity, PGRN^+/−^mice exhibited significantly increased perivascular AQP4 coverage relative to WT controls (Fig. 6E, F). We next examined pericyte association with the cortical vasculature by quantifying Caldesmon-positive coverage along CD31-positive microvessels. Caldesmon-positive perivascular coverage was modestly reduced in PGRN^+/−^ mice, although the difference did not reach statistical significance (Fig. 6G, H). These data therefore do not demonstrate substantial pericyte loss but are consistent with comparatively subtle changes in the perivascular organization. Finally, we assessed Iba1-positive microglial morphology as an indicator of neuroimmune alterations ^53^. IBA1-positive microglia from PGRN^+/−^ mice displayed significantly reduced process length and soma area relative to WT controls (Fig. 6I–K), indicating an altered microglial morphological state that accompanies vascular and astrocytic changes. Together, these results demonstrate that the effects of progranulin haploinsufficiency extend beyond the cerebral endothelium and are accompanied by coordinated remodeling of astrocytes, the astrocyte–vascular interface, and microglia, with more modest changes in Caldesmon-positive pericyte coverage.

**Figure 6.**
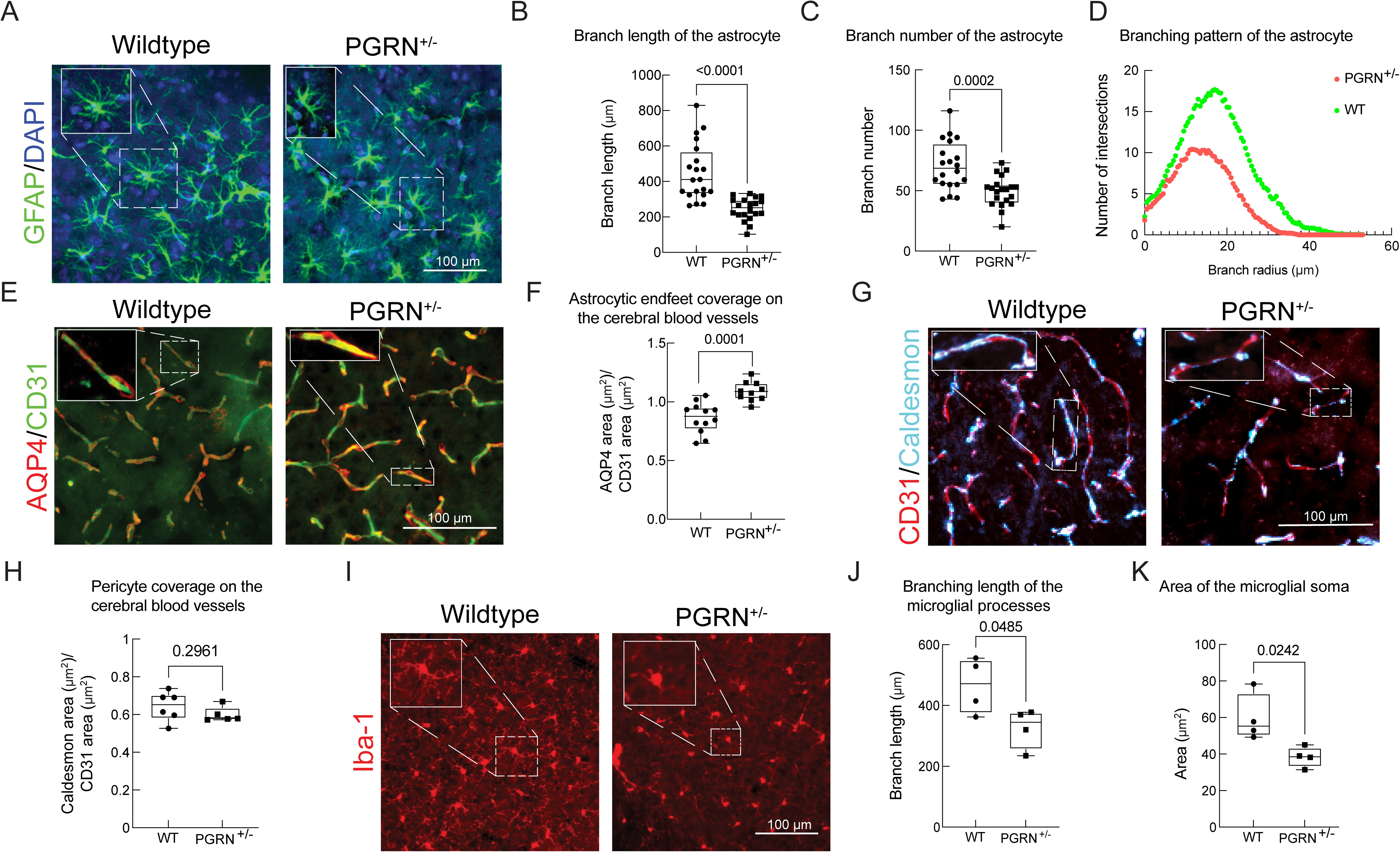
**Neurovascular unit cellular morphology in PGRN^+/−^ mice**. (A) Representative immunostaining of cortical astrocytes labeled with GFAP in WT and PGRN^+/−^ mice. Insets show individual astrocyte territories. Quantification of astrocyte branch length (B) and branch number (C) in WT and PGRN^+/−^ mice. (D) Sholl analysis of astrocytic branching complexity in WT and PGRN^+/−^ mice. (E) Representative immunostaining of astrocytic endfeet labeled with AQP4 and endothelial marker CD31 in WT and PGRN^+/−^ mice. (F) Quantification of AQP4-positive astrocytic endfoot coverage of CD31-labeled microvessels in WT and PGRN^+/−^ mice. (G) Representative immunostaining of pericytes labeled with Caldesmon and endothelial marker CD31 in WT and PGRN^+/−^ mice. (H) Quantification of pericyte coverage along CD31-labeled microvessels in WT and PGRN^+/−^ mice. (I) Representative immunostaining of cortical microglia labeled with Iba-1 in WT and PGRN^+/−^ mice. Insets show representative cellular morphology. Scale bar, 100 μm. Quantification of microglial cell body area (J) and branch length (K) in WT and PGRN^+/−^ mice. Data are presented as individual values with summary statistics. Statistical comparisons were performed as described in Methods

## DISCUSSION

Progranulin deficiency has been studied predominantly in the context of neuronal dysfunction, lysosomal biology, and microglial activation; however, its effects on the cerebral vasculature and neurovascular unit (NVU) remain comparatively understudied ^11,12,54^. To investigate these vascular alterations, we used heterozygous PGRN^+/−^ mice, which model the haploinsufficient state found in patients with FTD-GRN ^5,6,55,56^ and homozygous PGRN^−/−^ mice as a model of complete progranulin loss. By combining *in vivo* multiphoton imaging, cerebral microvessel transcriptional profiling, cross-species endothelial analysis, vascular protein measurements, blood–brain barrier (BBB) assessment, and NVU characterization, we identified a broad cerebrovascular phenotype associated with progranulin deficiency. These alterations included impaired cerebral perfusion, increased capillary stalling, endothelial immune activation, altered vascular maintenance signaling, BBB remodeling, and structural changes in multiple NVU cell types.

*In vivo* multiphoton imaging revealed impaired cerebral microvascular perfusion in progranulin-deficient mice, accompanied by increased capillary stalling. These microvascular changes in PGRN^+/−^ mice, the genotype that mirrors human FTD-GRN, indicate endothelial activation and early microvascular dysfunction in the haploinsufficient state, coincident with the behavioral and cognitive deficits documented in this mouse model ^15^. The finding that vessel diameter remained unchanged in PGRN^+/−^ mice argues against structural narrowing of vessels as the primary explanation, instead suggesting alterations in endothelial regulation, activation, and leukocyte-vascular interactions. Leukocyte adhesion within brain capillaries has been identified as a driver of stalled microvascular flow in Alzheimer’s disease mouse models, where even a small percentage of stalled capillaries produces disproportionate reductions in cortical blood flow ^38,57^. The elevated ICAM-1 expression in microvessels and increased leukocyte-capillary interactions observed here suggest a pro-inflammatory environment similar to that present in progranulin-deficient mice, as seen in Alzheimer’s disease mice ^38^. Broader evidence from the stroke literature further supports the vulnerability of the cerebral vasculature to progranulin status ^58^. PGRN has been shown to directly suppress TNF-α-induced ICAM-1 expression in brain microvascular endothelial cells and to attenuate neutrophil recruitment in ischemia-reperfusion injury models, while exogenous PGRN administration reduces infarct volume and limits hemorrhagic transformation in acute ischemic stroke ^59,60^. Chronic partial reduction of progranulin may therefore shift the cerebral endothelium toward a more adhesion-permissive state and increase susceptibility to capillary obstruction. Importantly, progressive reductions in CBF have been reported using arterial spin labeling MRI in presymptomatic FTD-GRN mutation carriers ^36^. Our findings provide a potential microvascular mechanism for this early blood flow reduction seen in FTD-GRN and FTD-MAPT patients and suggest that endothelial activation and capillary flow disturbances may emerge early in FTD-GRN.

Transcriptomic profiling of isolated cerebral microvessels revealed vascular reprogramming associated with progranulin haploinsufficiency. Differential gene expression analysis identified alterations in immune signaling, extracellular matrix organization, oxidative stress regulation, and vascular homeostatic pathways. PGRN^+/−^ microvessels showed downregulation of pathways related to angiogenesis, vascular development, and reactive oxygen species regulation, together with upregulation of extracellular matrix organization and tissue-remodeling programs. This combination is consistent with a shift away from an actively maintained vascular state toward a more dysfunctional or maladaptive endothelial phenotype ^20,61^. Also, a recent study has identified transcriptional changes suggesting premature vascular senescence in the arteries of a cardiorenal PGRN^−/−^ model ^62^.

Protein–protein interaction analysis further identified modules related to immune and lymphocyte signaling, chemokine-mediated leukocyte recruitment, antigen presentation, oxidative stress, vascular adhesion, and collagen organization. The enrichment of immune-associated pathways is particularly relevant given the established role of progranulin in suppressing inflammatory signaling in neurons and microglia ^12,13^. Our findings suggest that this regulatory role extends to the cerebral microvascular compartment and that partial progranulin loss may contribute to neuroinflammation through altered endothelial immune signaling, leukocyte recruitment, and vascular–immune interactions. The accompanying suppression of vascular-development and stress-response pathways indicates that endothelial immune activation occurs in parallel with impaired vascular homeostasis rather than as an isolated inflammatory response ^63^.

Cross-species pseudobulk transcriptomic comparison demonstrated partial conservation of these endothelial responses between progranulin-deficient mice and human FTD-GRN. Both PGRN^+/−^ and PGRN^−/−^ mice shared subsets of directionally concordant endothelial genes with the human disease signature, although substantial discordance was also present. The PGRN^−/−^ model reflects complete progranulin loss and produces a more severe inflammatory phenotype than the haploinsufficient state present in human mutation carriers. Concordance observed in the homozygous condition may therefore reflect convergence upon downstream inflammatory, metabolic, or stress-associated pathways rather than equivalence to the initiating human disease mechanism ^54,64–66^. Concordant genes shared between human FTD-GRN and the mouse models were enriched for processes including angiogenesis, endothelial migration, extracellular matrix organization, nitric oxide metabolism, endothelial proliferation, vascular permeability, and cell–cell junction organization. These pathways are directly relevant to vascular remodeling and BBB homeostasis ^9,20,67^. The cross-species analysis therefore does not indicate complete replication of human FTD-GRN by either mouse genotype, but it demonstrates that several central components of the vascular response to progranulin deficiency are conserved across species. This partial conservation strengthens the translational relevance of the endothelial alterations observed in the heterozygous model while also emphasizing that individual gene responses vary across species, anatomical regions, and disease stages. Protein-level analyses extended the transcriptomic findings and demonstrated that altered vascular programs were accompanied by structural rarefaction of the cortical vasculature. Angiogenesis-array profiling of PGRN^+/−^ cortical lysates identified significant reductions in NOV/CCN3 and Osteopontin. NOV/CCN3 is a matricellular CCN-family protein involved in endothelial survival, extracellular matrix regulation, and vascular remodeling ^68^. Its reduction is consistent with the transcriptional downregulation of angiogenesis-and vascular-development-associated pathways and supports impaired vascular maintenance as a feature of progranulin haploinsufficiency ^68^. The accompanying reduction in Osteopontin, a regulator of extracellular matrix signaling, inflammation, and tissue repair, further suggests disruption of vascular homeostatic signaling ^69,70^. VEGF-A levels were also reduced in PGRN^+/−^ cortical lysates, providing independent biochemical evidence of altered vascular trophic support. This finding is consistent with previous evidence that progranulin can promote VEGF expression and angiogenic responses ^71,72^. Importantly, these molecular changes were accompanied by a reduction in CD31-positive cortical vessels, consistent with the decreased vascular density. The convergence of transcriptomic, protein, histological, and intravital imaging findings therefore supports a model in which progranulin haploinsufficiency compromises vascular maintenance and contributes to cerebral microvascular rarefaction.

Additional reductions in PlGF-2, Proliferin, and Pentraxin-3 in PGRN^−/−^ mice suggest that complete progranulin loss produces broader disruption of vascular-associated protein expression. These findings should not, however, be interpreted as evidence that all vascular abnormalities progress monotonically with progranulin dose. Rather, the heterozygous and homozygous models appear to elicit overlapping yet distinct vascular responses.

The elevated ICAM-1 transcript signal detected in isolated cerebral microvessels was not fully reproduced by whole-cortical protein measurements. Gerrits et al. identified transcriptional upregulation of ICAM-1 and VCAM-1 in an inflammatory endothelial subcluster of the human FTD-GRN temporal cortex, supporting adhesion-molecule dysregulation as a feature of the disease endothelium ^18^. In the present study, soluble ICAM-1 and VCAM-1 concentrations were not significantly altered in PGRN^+/−^ cortical lysates. Several non-exclusive factors may explain this divergence. Isolated microvessel RNA preferentially captures endothelial and closely associated vascular transcripts, whereas cortical ELISA measures soluble protein derived from multiple vascular and nonvascular cell types. In addition, transcript abundance, endothelial surface protein levels, and soluble adhesion molecule concentrations represent distinct molecular pools.

Following the demonstration of impaired vascular maintenance and cortical microvascular rarefaction, we examined whether progranulin deficiency altered BBB organization. Claudin-5 coverage along cortical vessels was significantly reduced in PGRN^+/−^ mice, while Occludin vessel association and total Claudin-5 and Occludin protein abundance showed nonsignificant downward trends. These findings indicate altered tight junction organization rather than a complete loss of junctional proteins ^20,73^. Despite these molecular and structural alterations, neither endogenous IgG immunoreactivity nor sodium fluorescein accumulation was significantly increased. The absence of overt leakage despite altered junctional organization is biologically plausible. Claudin-5 and Occludin can redistribute away from inter-endothelial junctions or undergo partial downregulation before permeability changes become detectable using bulk tracer assays, a pattern described during early endothelial stress and neurodegeneration ^21,74^. However, two studies, Cheemala et al. and Aragón-González et al., showed that an ALS/FTD-linked TDP-43 mutation and C9orf72, both models of FTD, respectively, impair endothelial barrier function, while Pan et al. showed broader BBB alterations in FTD, which together establish that cerebrovascular dysfunction is an emerging feature of the FTD-spectrum ^19,20,75^. Our findings may represent an earlier state in FTD-GRN, in which tight-junction organization, transporter expression, and endothelial signaling are altered before measurable basal leakage develops. Downregulation of solute carrier pathways further suggests that transcellular transport functions may be compromised in ways not captured by a fluorescein permeability assay ^51^. These observations are consistent with the concept that BBB dysfunction exists along a continuum, ranging from altered endothelial transport and signaling to frank structural breakdown ^51,76,77^. The BBB-associated PPI modules further support complex endothelial remodeling. The extracellular matrix and barrier-disruption module contained increased Mmp14 and Timp1 expression together with reduced Mmp12 and Pappa expression. This mixed pattern indicates reorganization of protease and protease-inhibitor balance rather than uniform matrix degradation. Increased Mmp14 may promote pericellular matrix remodeling, while increased Timp1 may represent a simultaneous compensatory attempt to restrain excessive proteolysis ^45^. The reductions in Mmp12 and Pappa further indicate that distinct extracellular proteolytic pathways are differentially affected. The innate immune module showed increased Ager and Pnpla3 expression together with reduced Cd36 and Emp1 expression, supporting dysregulation of inflammatory, lipid-associated, scavenger-receptor, and membrane-signaling pathways ^48,49^. The coagulation and blood-derived signaling module was characterized by increased Ambp, Pan, Serpinc1, and Tfpi2 together with reduced Klkb1, suggesting remodeling of anticoagulant, protease-inhibitory, and circulating protein-associated pathways ^50^. Collectively, these changes indicate that BBB dysfunction in progranulin deficiency is characterized by broad endothelial and molecular reprogramming rather than overt barrier rupture.

The cerebral vasculature may nevertheless be more vulnerable to physiological or pathological challenge in the setting of progranulin deficiency. PGRN has been shown to limit BBB disruption and regulate vascular permeability through VEGF-associated mechanisms following ischemic injury ^60^. Thus, preservation of basal tracer exclusion does not necessarily imply normal barrier resilience. The progranulin-haploinsufficient vasculature may possess reduced reserve capacity and may be disproportionately susceptible to inflammation, ischemia, neuronal activation, or aging.

Endothelial and BBB alterations were accompanied by structural remodeling of multiple non-endothelial components of the NVU. Morphological changes in astrocytes and microglia are consistent with previous findings in progranulin-deficient mouse models and postmortem FTD-GRN tissue ^18,54,65^. GFAP-positive astrocytes in PGRN^+/−^ mice exhibited reduced branch length, branch number, and Sholl complexity, indicating simplified astrocytic arborization. In contrast, AQP4-positive coverage along CD31-positive vessels was increased, suggesting redistribution of astrocytic organization toward the vascular interface rather than a uniform loss of astrocytic structure.

Astrocytic endfeet ensheathe cerebral vessels and play central roles in BBB maintenance, water balance, ion homeostasis, and neurovascular coupling. Increased perivascular AQP4 coverage may therefore reflect a reactive or compensatory response to endothelial dysfunction, although increased coverage should not automatically be interpreted as improved function ^18,44^. Notably, Gerrits et al. independently reported increased perivascular AQP4 coverage in the frontal cortex of postmortem FTD-GRN tissue, suggesting that reactive astrocytic remodeling at the glia-vascular interface is a conserved feature across the disease trajectory ^18^. Caldesmon-positive pericyte coverage showed a modest but nonsignificant reduction in PGRN^+/−^ mice. The current data therefore do not establish substantial pericyte loss. Nevertheless, the direction of change is consistent with the reported reduction in PDGFRB-positive pericyte coverage in human FTD-GRN cortex and supports further investigation using additional pericyte markers and functional assays.^18,78^.

Microglia in PGRN^+/−^ mice exhibited reduced process length and soma area. These findings demonstrate altered microglial morphology but do not, by themselves, define a specific activated or inflammatory state. Microglial morphology is multidimensional and context-dependent, and similar structural changes can occur across several reactive, degenerative, or transitional phenotypes. The microglial findings are therefore interpreted as evidence of structural remodeling accompanying vascular and astrocytic alterations. Together, the astrocytic, pericytic, and microglial changes demonstrate that the consequences of progranulin haploinsufficiency extend beyond the endothelium and involve broader reorganization of the NVU.

Several limitations of the present study should be considered. The analyses presented here are cross-sectional, conducted in 10–12-month-old mice at a single time point; consequently, the temporal sequence of neurovascular changes, whether microvascular perfusion deficits precede BBB remodeling, or whether NVU structural changes arise before or after endothelial transcriptional reprogramming, cannot be determined from the current dataset. Longitudinal studies will be required to establish the order of events and determine whether these changes progress with age or correlate with the behavioral changes that emerge in this model ^15^, a question important for establishing whether cerebrovascular dysfunction mechanistically contributes to behavioral changes in progranulin deficiency. Microvessel RNA sequencing libraries were generated from pooled samples due to limited RNA yield per animal, and transcriptomic results are therefore interpreted as genotype-associated expression patterns rather than estimates of individual biological variability; however, key transcriptional observations were supported by orthogonal approaches. BBB permeability was assessed under baseline conditions. Progranulin deficiency may increase vascular vulnerability under physiological stress, neuronal activation, or disease progression in ways not captured by a resting-state assay ^76,77^. Evidence from ischemia models demonstrating that PGRN loss worsens cerebrovascular injury outcomes ^59,60^ suggests that the haploinsufficient vasculature characterized here may be disproportionately susceptible to such challenges. While microvessel isolation enriches for vascular transcripts, contributions from closely associated perivascular cell types such as astrocytic endfeet and pericytes are likely to occur.

In summary, progranulin haploinsufficiency is associated with neurovascular remodeling, encompassing impaired microvascular perfusion with endothelial activation, transcriptional reprogramming of the cerebral endothelium, reduced angiogenic and vascular homeostatic signaling, BBB reorganization, and structural alterations across NVU cell types. Vascular neuroinflammation, microvascular stalling, immune pathway enrichment in the endothelial transcriptome, and structural microglial remodeling emerge as central features and may represent key mechanisms through which progranulin haploinsufficiency disrupts neurovascular homeostasis. These findings are consistent with, and provide cellular-level mechanistic context for, the CBF reductions observed in presymptomatic GRN mutation carriers ^36^ and the neurovascular transcriptional alterations identified in end-stage FTD-GRN postmortem tissue ^18^, linking observations across the human disease trajectory to a tractable *in vivo* model of haploinsufficiency. Increasing recognition that endothelial dysfunction is a shared feature of the broader FTD-ALS disease spectrum, including disease caused by TDP-43 and C9orf72 mutations ^19,20,61^, underscores the importance of integrating cerebrovascular biology into disease frameworks and therapeutic avenues to increase PGRN ^32,33,79,80^.

## MATERIAL and METHODS

### Animals

The PGRN-deficient mice were ordered from Jackson Labs (#:013175) and maintained at the University of Miami ^15^. The model has a loss of the GRN and shows a progressive frontotemporal dementia. In this study, we primarily used heterozygous PGRN^+/-^ mice (reflecting more FTD-GRN), and for some experiments, we included homozygous PGRN^-/-^mice. In this study, craniotomies for *in vivo* imaging were performed on 8 PGRN^+/-^, 7 PGRN^-/-^, and 6 wildtype (WT) controls aged 11 to 13 months, including both males and females. In total, we used 31 wildtype mice, 25 PGRN^+/-^, and 30 PGRN^-/-^mice of both sexes. Mouse brains were harvested after intracardiac perfusion. Animal colonies were maintained in the Department of Biology at the University of Miami. All animal procedures were conducted in accordance with the National Institutes of Health (NIH) Guide for the Care and Use of Laboratory Animals and approved by the institutional animal care and use committee (IACUC) at the University of Miami.

**Table 1:** Animal number.

| Experiment name | Wildtype mice number | PGRN<sup>+/-</sup> mice number | PGRN<sup>-/-</sup> mice number |
| --- | --- | --- | --- |
| In-vivo Imaging | 6 | 7 | 7 |
| Immunostaining | 8 | 6 | 6 |
| Western Blot,<br>ELISA,<br>Angiogenesis array | 8 | 7 | 8 |
| Sodium-Fluorescein assay | 9 | 5 | 9 |

### Immunostaining

Mouse brains were sectioned sagittally at 20 microns for immunostaining experiments ^81^. Following removal from cryoprotectant, sections were washed four times with phosphate-buffered saline (PBS). Pretreatment solutions were applied based on the target antigen: sodium borate for Iba-1, and sodium citrate for Caldesmon and GFAP. Tris/EDTA was used for Claudin-5 and Occludin ^82^. Sodium borate pretreatment was performed for 30 minutes at room temperature, while sodium citrate pretreatment was carried out at 95 °C for 25 minutes with constant agitation. After pretreatment, samples were washed three to five times with PBS and then incubated in a 5% goat serum-blocking solution for 1 hour at room temperature. Primary antibodies were applied, and samples were incubated overnight at 4 °C with gentle agitation on a rocker. The following day, sections were washed three to four times with PBS before incubation in secondary antibody solutions for 90 minutes at room temperature with gentle agitation on a rocker. Finally, samples were washed three times with PBS, mounted with DAPI-containing mounting media, and allowed to dry at room temperature. Imaging was performed using a Zeiss LSM 880 confocal microscope. Image acquisition focused exclusively on the cerebral cortex in sagittal sections, using 63x, 20x, and 10x objective lenses.

**Table 2:** Antibodies.

| Primary antibody | Primary antibody concentration | Pretreatment | Catalogue number |
| --- | --- | --- | --- |
| Iba-1 | 1/150 | 0.1% sodium borate buffer at room temperature for 30 minutes | Anti-Iba1 antibody [EPR16588] (ab178846) |
| GFAP | 1/250 | Citrate buffer at 95°C for 25 minutes | Anti-GFAP antibody (ab4674) |
| Aqp-4 | 1/1000 | Citrate buffer at 95°C for 25 minutes | Aquaporin 4 Antibody (PA5-53234) |
| CD-31 | 1/20 | Citrate buffer at 95°C for 25 minutes | Anti-CD31 (Ms, Sw) from Rat (SZ31) |
| Caldesmon-1 | 1/200 | Citrate buffer at 95°C for 20 minutes | Caldesmon-1 (D5C8D) XP® Rabbit mAb #12503 |
| Lectin | 1/250 | Not required | Invitrogen™<br>Lycopersicon<br>Esculentum (Tomato)<br>Lectin (LEL, TL),<br>fluorescein (FITC) |
| Claudin-5 | 1/20 | Tris/EDTA buffer at 95°C for 20<br>minutes | Claudin 5 Monoclonal<br>Antibody (4C3C2) |
| Occludin | 1/200 | Tris/EDTA buffer at 95°C for 20<br>minutes | Occludin Monoclonal<br>Antibody (OC-3F10) |

### Image analysis

Immunostained slides were imaged using a Zeiss laser scanning confocal microscope (LSM) 880. Image analysis pipelines were customized according to the specific research question. **Blood Vessel Number:** Lectin-stained slides were imaged, and the resulting.czi files were opened in ImageJ software ^83^. Individual blood vessel segments were selected using the polygon selection tool, added to the ROI manager, and various parameters were measured using the Measure tool. The output was saved in an Excel file. For each mouse, three images were analyzed, and the values were averaged. **Pericyte Coverage of Cerebral Blood Vessels:** Caldesmon and CD31 markers were used for immunostaining. Images were opened in ImageJ. First, CD31+ blood vessel segments were selected using the polygon selection tool, added to the ROI manager, and their total area was measured. Subsequently, Caldesmon+ blood vessel segments were measured similarly, but only those segments previously identified as CD31+ were selected for analysis, ensuring that only the area of Caldesmon fluorescence co-localizing with CD31 was considered. **Astrocytic Endfeet Coverage of Cerebral Blood Vessels:** The same methodology as pericyte coverage was followed to measure astrocytic endfeet coverage on cerebral blood vessels, using Aqp-4 and CD31 as markers (Supplementary Fig. 3A–B). **Astrocyte Branching Pattern Analysis:** Alterations in astrocyte branching patterns were assessed using the Simple Neurite Tracer (SNT) plugin in ImageJ ^84^. Briefly, GFAP-and DAPI-immunostained images were opened, and individual astrocytes positive for DAPI were selected. The SNT plugin was initiated, and the center of the DAPI nucleus was marked as the starting point. Main astrocyte branches were traced from this center, followed by tracing of side branches originating from the main branches. After tracing all branches from a single astrocyte, the tracings were saved and used to calculate branch length, number, and other parameters. Additionally, Sholl analysis was performed by drawing concentric circles from the astrocyte’s center point and measuring the number of intersections between astrocyte branches and circles of a particular diameter, which aids in determining branching pattern alterations ^85,86^ (Supplementary Fig. 4). **Microglial Morphology Analysis:** To assess shifts in microglial morphology, Iba-1 immunostained mouse brain sections were imaged and opened in ImageJ and the following methodology is followed ^87^. The single Iba-1 channel was selected. A randomization and grid-creation script was used to divide the image into multiple grids, each approximately the size of an individual microglial cell. Five microglia were randomly selected from these grids, and three of these five were chosen based on the clarity of their branches and the clear presence of the microglial cell body. Each of these three microglial images was analyzed individually. Duplicate images were created and modified with various filters to enhance visualization of specific morphological features, but the original single-microglia picture was used for downstream analysis. First, the image was converted to binary, and the ImageJ brush tool was used to remove extraneous branches from other microglia, artifacts, and noise. The image then underwent multiple filters, including despeckle, denoise, and remove outliers. Subsequently, the image was skeletonized, duplicated, and the skeleton was analyzed using the Analyze Skeleton tool in ImageJ ^88^. Additionally, the cell body area of each microglia was measured using the polygon selection tool. Results were stored in an Excel file. This methodology was applied to three microglia per image, and the corresponding parameter values were averaged per image (Supplementary Fig. 5). **Tight Junction Protein Colocalization (Blood-Brain Barrier Integrity):** Claudin-5 and Occludin, along with Lectin, were stained to assess alterations in blood-brain barrier integrity. Images were opened in ImageJ, fluorescent channels were separated, and the JACOP plugin in ImageJ was used to measure colocalization of Claudin-5 with Lectin and Occludin with Lectin ^89^. Pearson’s correlation coefficients from these analyses were stored in an Excel file.

### Microvessel isolation and RNA sequencing

After collecting the brains from the mice, they are chopped into fine pieces in Qiazol ^90,91^. Using stainless steel beads, the brain pieces are further homogenized. The homogenate is then centrifuged at 12000 g for 10 minutes at 4 °C. The solution is separated into three distinct layers: fat (top), RNA (middle), and pellet (bottom). The middle layer containing the RNA is taken out separately, and chloroform is added, mixed, and centrifuged at 10000 g at 4°C for 15 minutes. The upper aqueous phase is then mixed with 70% ethanol in a 1:1 ratio. RNA is isolated with the modified protocol of the RNeasy Mini Kit. The solution is passed through the spin column, centrifuged, and then sequentially washed with RW1 and RPE buffers. Finally, the RNA is eluted in RNase-free water, its quality is assessed on a NanoDrop, and it is sent for sequencing.

### Human Endothelial Pseudobulk Transcriptomic Analysis

Publicly available single-nucleus RNA sequencing (snRNA-seq) data (GSE163122) from the cerebral cortex of frontotemporal dementia (FTD-GRN) patients and non-demented controls generated by Gerrits et al. were obtained and reanalyzed in R using the Seurat and edgeR packages ^18,92,93^. Raw count matrices were imported using the Read10X function. For exploratory visualization and marker validation, nuclei were randomly downsampled to 20,000 cells per sample prior to normalization. Data were normalized using NormalizeData, highly variable genes were identified with FindVariableFeatures, and scaled expression matrices were generated with ScaleData. Dimensionality reduction was performed using principal component analysis (RunPCA), followed by graph-based clustering (FindNeighbors, FindClusters) and UMAP embedding (RunUMAP). Endothelial nuclei were identified based on expression of canonical marker genes including CLDN5, PECAM1, FLT1, KDR, RAMP2, and ESAM. Nuclei were classified as endothelial if they expressed at least one of these markers and lacked expression of immune lineage markers (PTPRC or LST1), a criterion applied to minimize inclusion of contaminating leukocytes. Endothelial nuclei were assigned to arterial, capillary and venous segments using canonical zonation markers ^78,94,95^, and differential expression between FTD-GRN and control capillary endothelial nuclei (1,981 FTD-GRN and 2,104 control nuclei) was tested with MAST (Seurat, FindMarkers), identifying 1,223 genes at FDR < 0.05. These genes were used for the cross-species comparison. For pseudobulk analysis, raw count matrices were reprocessed without downsampling to preserve the full cellular complement. Following marker-based filtering, endothelial nuclei were retained and raw counts were aggregated by summing across all qualifying nuclei within each anatomical region to generate region-level pseudobulk profiles. Region-level profiles from the same donor were subsequently summed to produce a single donor-level endothelial pseudobulk expression vector. This aggregation strategy yields sample-level count data appropriate for bulk RNA-seq statistical frameworks and mitigates inflated false-positive rates associated with single-cell-level differential expression testing ^96^.

Donor-level pseudobulk profiles were generated for four control donors (C2, C3, C4, C5) and ten FTD-GRN donors (P1, P2, P3, P4, P5, P8, P11, P12, P16, P20). Following marker-based filtering, 6,352 endothelial nuclei were retained for pseudobulk aggregation and assigned to arterial (n = 915), capillary (n = 4,085) and venous (n = 1,352) segments. Differential expression between FTD-GRN and control endothelial pseudobulk profiles was assessed using the edgeR package. Genes expressed at low levels across samples were removed using the filterByExpr function with default parameters. Library sizes were normalized using the trimmed mean of M-values (TMM) method to account for compositional differences between samples. Dispersion parameters were estimated within a generalized linear model framework, and differential expression was tested using quasi-likelihood F-tests (glmQLFit, glmQLFTest) with diagnostic group as the primary covariate ^97^. Results are reported as log₂ fold changes with associated false discovery rate (FDR)-adjusted p-values.

For quality control and visualization, log-transformed counts per million (logCPM) were computed using the cpm function, and principal component analysis of the logCPM matrix was used to assess sample-level clustering. Volcano plots were generated from the quasi-likelihood model output.

### Cross-Species Endothelial Transcriptomic Comparison and Pathway Enrichment

To evaluate the translational relevance of endothelial transcriptional changes observed in progranulin-deficient mice, cross-species comparisons were performed between mouse cerebral microvessel differential expression datasets and human endothelial pseudobulk differential expression results derived from the Gerrits et al. dataset. Mouse differential expression datasets consisted of PGRN⁺/⁻ vs wild-type and PGRN⁻/⁻ vs wild-type cerebral microvessel RNA-seq comparisons representing heterozygous and homozygous progranulin-deficiency models. Prior to integration, gene symbols were standardized by converting all human and mouse identifiers to uppercase and removing entries with missing or duplicated symbols. Human differential expression tables were reduced to gene symbol, log₂ fold change, and FDR, while mouse heterozygous and homozygous tables were formatted to retain gene symbol, log₂ fold change, and adjusted p-value.

Ortholog mapping between mouse and human genes was performed using the babelgene package ^98^. genes from both datasets were pooled and mapped to their human orthologs, and the resulting mapping table was filtered to retain non-missing, one-to-one ortholog pairs.

Mouse differential expression values were then joined to the corresponding human ortholog entries. For each comparison, heterozygous mouse versus human and homozygous mouse versus human overlapping orthologous genes were identified and assessed for directional concordance. Genes were classified as showing *same-direction regulation* when the sign of the mouse log₂ fold change matched that of the human log₂ fold change, and *opposite-direction regulation* otherwise. Pearson correlations were calculated between mouse and human log₂ fold changes across all overlapping orthologs. Scatterplots were generated in ggplot2, with axes demarcated at zero, points colored by directional concordance, and selected gene labels rendered using ggrepel^99^.

Genes reaching statistical significance in both species were identified using an adjusted p-value threshold of 0.05 in the mouse datasets and an FDR threshold of 0.05 in the human capillary dataset. Genes shared across human, heterozygous mouse, and homozygous mouse comparisons were additionally identified and ranked by human differential expression significance (FDR), prioritizing genes most confidently dysregulated in human FTD endothelial cells while evaluating whether these disease-associated transcriptional changes were recapitulated directionally in mouse models. The 45 capillary genes with the strongest separation between FTD-GRN and control donors were visualized as a heatmap of row-wise z-scores of per-donor pseudobulk logCPM using pheatmap, with rows and columns ordered by hierarchical clustering (Euclidean distance, complete linkage) ^100^.

For pathway enrichment analysis, cross-species overlap tables were filtered to retain genes displaying same-direction regulation between species, under the rationale that conserved directional dysregulation is more likely to reflect shared pathobiological mechanisms. Mouse gene symbols from these concordant overlap sets were converted to human Entrez identifiers using clusterProfiler and the org.Hs.eg.db annotation database ^101^. Gene Ontology biological process enrichment was performed using clusterProfiler (enrichGO, ont = BP) with Benjamini–Hochberg multiple-testing correction, as previously described ^78,102,103^.

To enable full pathway ranking prior to visualization, permissive significance thresholds were applied (pvalueCutoff = 1, qvalueCutoff = 1). Enriched terms were ranked and visualized by p-value. Enriched pathways are displayed as dot plots with point size proportional to gene count and point color scaled to −log₁₀(p-value), with pathway labels wrapped for readability. Enriched terms were ranked by p-value, and the terms displayed in Fig. 3D, E were restricted to vascular and angiogenesis-associated biological processes (angiogenesis and sprouting, endothelial migration and proliferation, extracellular matrix organization, cell adhesion and junctions, vascular permeability and homeostasis). Complete enrichment tables are provided in Supplementary Table 2. All ortholog mapping tables, cross-species overlap tables, pathway enrichment results, and figures were written to disk for downstream manuscript assembly.

### ELISA and Angiogenesis Array

Cortical tissue was homogenized in a custom lysis buffer consisting of 20 mM Tris base, 150 mM NaCl, 1 mM EDTA, 0.1% Triton X-100, and 5% glycerol supplemented with a protease inhibitor cocktail. Homogenates were centrifuged to remove insoluble debris, and supernatants were collected and stored at −80°C until use. Total protein concentration in each homogenate was determined by the BCA assay to normalize analyte concentrations across samples. Levels of soluble ICAM-1, VCAM-1, and VEGF-A were quantified using commercially available ELISA kits: the Quantikine Mouse ICAM-1/CD54 Immunoassay (R&D Systems, Cat# MIC100), the Mouse VCAM-1 SimpleStep ELISA Kit (Abcam, Cat# ab201278), the Mouse VEGF-A ELISA Kit (Invitrogen, Cat# BMS619-2), and the Proteome Profiler Mouse Angiogenesis Array Kit (R&D Systems, Cat# ARY015), respectively. All assays were performed on cortical tissue homogenates according to each manufacturer’s instructions, with sample dilutions optimized empirically to ensure readings fell within the linear dynamic range of each assay. All samples were assayed in duplicate, and mean absorbance values were interpolated against a freshly generated standard curve for each plate using a four-parameter logistic (4-PL) curve fit. Final analyte concentrations were normalized to total protein content and expressed as pg/mg total protein. BioTek Synergy H1 Hybrid Multi-mode Microplate Reader was used to detect the absorbance in these ELISAs

### Western blotting

One hemisphere of the mouse cerebral cortex was dissected and homogenized in 500 µL of protein extraction buffer consisting of RIPA buffer and PBS (1:25) supplemented with protease inhibitor cocktail (Sigma-Aldrich). Tissue was disrupted by 20–30 gentle strokes in a Dounce homogenizer, taking care to avoid bubble formation, followed by brief sonication. Lysates were then centrifuged at 15,000 × *g* for 30 minutes at 4°C, and the resulting supernatant was aliquoted and stored at –80°C until use. Protein samples (20 µg) were resolved on a 15% SDS-PAGE gel at 75 V initially, then at 90 V until the dye front reached the end of the gel. Proteins were transferred to a PVDF membrane by wet transfer at a constant 80 V for 90 minutes. Membranes were blocked in 3% BSA in PBS-T for 1 hour at room temperature, then incubated overnight at 4°C with primary antibodies diluted in blocking solution: anti-Claudin-5 (4C3C2, Thermo Fisher 35-2500; 1:500), anti-Occludin (OC-3F10, Thermo Fisher 33-1500; 1:500), and anti-β-Actin (mAbcam 8226, ab8226; 1:5,000). Following primary incubation, membranes were probed with IRDye 800CW-conjugated secondary antibodies (1:5,000 in blocking solution). The fluorescent signal was detected at 700 nm and 800 nm simultaneously using an Odyssey imaging system, enabling co-visualization of the molecular weight ladder and target proteins.

### Cranial Window Surgery

Surgery was done as in ^104^. Briefly, the mice were anesthetized with 3% isoflurane using a Low-Flow Electronic Vaporizer (Kent Scientific) and maintained on 1.2% to 1.5% isoflurane in air delivered using a stereotaxic nose cone mask. Prior to surgery, mice were administered subcutaneous injections of atropine (Med-Pharmex, 54925-063-10; 0.005 mg/100 g) to reduce pulmonary secretions, dexamethasone (Phoenix Pharm, 07-808-8194; 0.025 mg/100 g) to limit postoperative inflammation, and ketoprofen (Zoetis; 0.5 mg/100 g) for analgesia and additional anti-inflammatory effects. A local anesthetic nerve block was applied at the incision site using 0.1 mL of 0.125% bupivacaine (Hospira). It is further supplemented with 5% (w/v) glucose in physiological saline (5 ml per kg of mouse) every hour. Body temperature is measured and maintained at 37.5 °C using a heating blanket. Heart rates are monitored using a custom-made MATLAB-based device. Eyes are protected by a covering of veterinary eye ointment. All surgical areas are shaved and cleaned with povidone-iodine, followed by swabbing with 70 % (v/v) alcohol. A 6 mm craniotomy is prepared over the parietal cortex, while the dura is left intact. A glass coverslip will be glued to the skull to close the craniotomy. For post-surgical care, subcutaneous injections of dexamethasone (0.025 mg/100 g) and ketoprofen (0.5 mg/100 g) were given for 2 days.

### Two-photon microscope (2PEF) imaging

Imaging was initiated four weeks after surgery. For *in vivo* two-photon (2PEF) microscopy, mice were anesthetized with 3% isoflurane for induction and maintained at 1.2–1.5% via a nose cone. Body temperature was continuously monitored and regulated between 36.5 and 37.5 °C throughout imaging sessions. Prior to imaging, mice received retro-orbital injections of Texas Red dextran (50 µL, 2%, MW 70,000 kDa; Thermo Fisher Scientific) to visualize the microvasculature, Rhodamine 6G (0.1 mL, 1 mg/mL in 0.9% saline; Millipore Sigma) to label leukocytes, and Hoechst 33342 (50 µL, 4.8 mg/mL in 0.9% saline; Thermo Fisher Scientific) to label platelets and distinguish them from leukocytes. Three-dimensional images of the cortical vasculature were acquired, and red blood cell (RBC) speeds, diameters, volumetric flow, and flux were measured in capillaries and penetrating arterioles. Imaging was performed using an InSight® X3+ laser (Spectra-Physics) tuned to 830 nm with 120 fs pulses. Galvanometric scanners operating at 1 frame per second directed the laser through a 20× water-immersion objective lens (NA 1.0; Carl Zeiss Microscopy). Fluorescence emission was detected with a four-channel system on a Bergamo II microscope (Thorlabs). The optical pathway included a 562 nm long-pass dichroic, followed by additional long-pass dichroics at 495 and 635 nm, and bandpass filters of 447/60, 525/50, 607/70, and 647 LP. Image acquisition and laser scanning were controlled using Thorlabs software. Z-stack images were collected at 1 µm axial intervals over the somatosensory cortex, reaching depths of 300–600 µm to visualize the cortical vasculature. Capillary stalling was quantified using ImageJ, and blood flow was measured with custom-built MATLAB code ^38^.

To quantify CBF in cortical penetrating arterioles, vessel diameter was measured from image stacks, and capillary red blood cell (RBC) velocity was obtained from line-scan measurements ^104,105^. The same capillaries were imaged repeatedly, and blood flow was measured in six to eight penetrating arterioles per mouse and 15-20 capillaries. All analyses were conducted in a blinded manner with respect to genotype, age, treatment, and imaging time point. RBC flux was derived from time traces of the capillary line scans. RBC peaks within these sequences were identified in MATLAB using the findpeak function.

### Sodium Fluorescein assay

Stock solutions of fluorescein sodium salt (F6377, Sigma-Aldrich) were prepared in 0.1 N NaOH, and standard solutions of 100, 50, 25, 12.5, 6.25, 3.12, 1.6, 0.8, 0.4, 0.2, 0.1, 0.05, 0.025, and 0.0125 μM were prepared in PBS. Mice were anesthetized, and fluorescein sodium salt diluted in PBS was injected retro-orbitally at a dose of 120 mg/kg. The dye was allowed to circulate for 15 min while the animal was monitored carefully under continuous anesthesia. After 15 min, the mice were euthanized and the brain was collected quickly, weighed, and homogenized in 500 μl PBS, to which an equal volume of 20% TCA was added. The solution was kept on ice in the dark for 15 min and then centrifuged at 15,000 g at 4 °C for 30 min. The supernatant was collected into a separate tube, and its pH was measured and adjusted with pH strips until it was between 7.4 and 8.0. The supernatant was then loaded into a 96-well plate in the dark and measured at 485 nm excitation and 530 nm emission in a microplate reader alongside the standards. The resulting fluorescein concentration was normalized to the brain weight of each animal.

### Statistics

All statistical analyses were performed using GraphPad Prism (version 10.1.1; GraphPad Software, San Diego, CA, USA). Data are presented as mean ± SEM unless otherwise indicated. For comparisons between two groups, two-tailed Mann–Whitney tests were used for nonparametric data and unpaired two-tailed Student’s t-tests were used for normally distributed data. When the variance between groups was unequal, Welch’s correction was applied. For comparisons involving more than two groups, a one-way ANOVA was used, followed by Holm–Šídák or Tukey’s multiple comparisons tests, for parametric data. When data did not meet the assumption of normality, Kruskal–Wallis tests, followed by Dunn’s multiple comparisons tests, were performed. For repeated-measures or longitudinal datasets, two-way ANOVA or mixed-effects models (with the Geisser–Greenhouse correction where appropriate) were used to account for within-subject variability and missing values. Multiple comparisons were corrected within each analysis as indicated. Statistical significance was defined as p < 0.05. Exact p-values, statistical tests, and sample sizes (n) are provided in the corresponding figure legends.

## Acknowledgments

We would like to thank Drs Chris Schaffer, Nozomi Nishimura, and Fenghua Hu for providing feedback and the mouse line. Thank you to all lab members for critical reading the manuscript.

## Declaration of conflicting interests

The authors declared no potential conflicts of interest with respect to the research, authorship, and/or publication of this article.

## Funding

The study was supported by the National Institute of Neurological Disorders and Stroke (NINDS) (NS141704 to S.T. and NS141137 to O.B), National Institute of Aging (AG075798 and AG082193 O.B), Alzheimer’s Association (AARG-22-974437 O.B.), Florida Department of Health (23A12 O.B), American Heart Association (American Student Scholarships in Cardiovascular Disease and Stroke N.N.B), Diversity supplement NIA (AG075798-01A1S1 (S.A.F.), and a McKnight Doctoral Fellowship from the Florida Education Fund (G.R.M.).

## Author Contributions

S.C. and O.B. contributed to overall project design. S.C., M.W., S.A.F., Z.T., C.S., G.R., D.L., N.N.B., A.C., P.P., S.T., and O.B. performed experiments and analyzed data. S.C. and R.R. performed cross species analysis. S.C. and O.B. wrote the manuscript and all authors edited the manuscript. All authors read and approved the final manuscript.

## Notes

### Competing Interest Statement

The authors have declared no competing interest.

