## SUPPLEMENTARY MATERIAL for "Progranulin haploinsufficiency remodels the cerebral microvasculature and neurovascular unit"

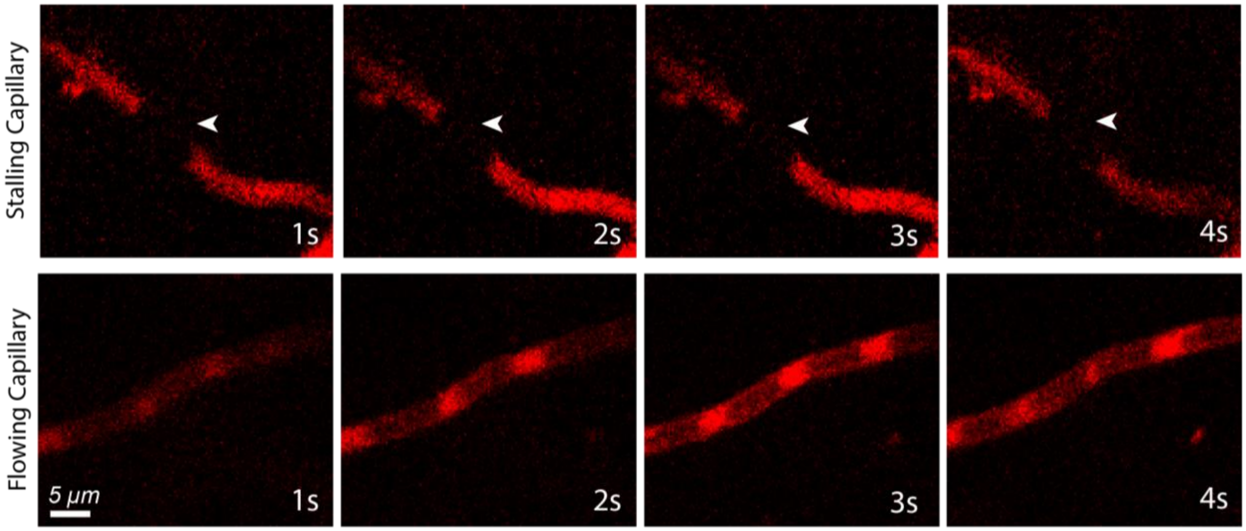


**Supplementary Figure 1. Example of a flowing and stalled capillary.** Individual brain capillaries were scored as stalled (top panel) or flowing (bottom panel) based on the motion of unlabeled blood cells (black) within the fluorescently labeled blood plasma (red).

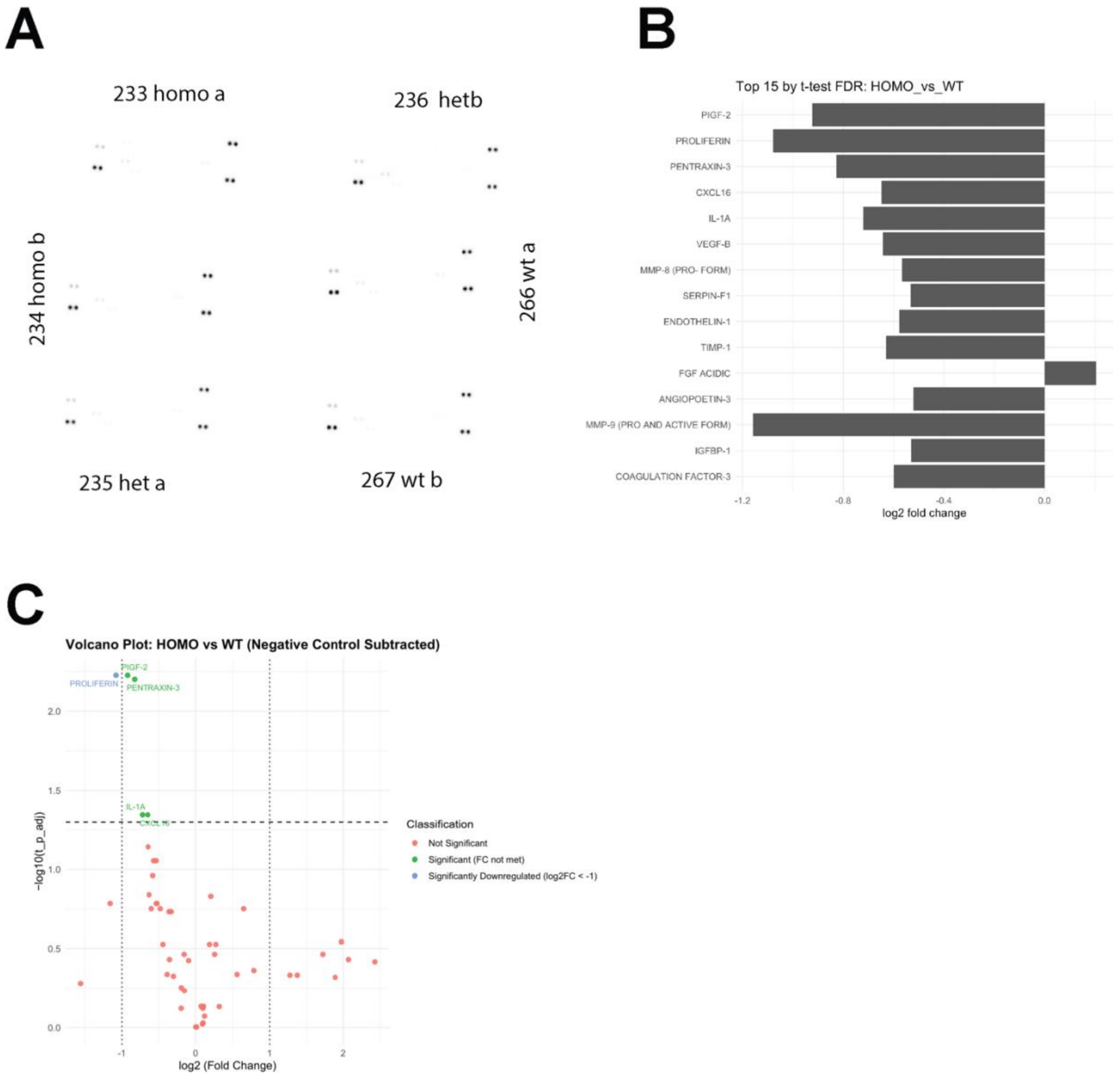


**Supplementary Figure 2. Raw angiogenesis array membranes and pairwise PGRN$^{-/-}$ versus WT angiogenic protein analysis. (A)** Raw angiogenesis array membranes from all six samples: two homozygous PGRN$^{-/-}$ (233a, 234b), two heterozygous PGRN$^{+/-}$ (235a, 236b), and two wild-type (266a, 267b) mice. Dot intensity reflects relative protein abundance across the 55-target panel and dot pairs are technical duplicates. **(B)** Top 15 angiogenesis-associated proteins ranked by t-test FDR in the PGRN$^{-/-}$ versus WT comparison, expressed as $\log_2$ fold change relative to WT; PIGF-2, PROLIFERIN, and

PENTRAXIN-3 are the top-ranked targets by FDR. **(C)** Volcano plot of pairwise PGRN$^{-/-}$ versus WT differential protein expression after negative-control subtraction. Horizontal dashed line, FDR threshold; vertical dashed lines, $|\log_2 FC| = 1$. Points are classified as not significant (red), significant by FDR but not fold change (green), or significantly downregulated meeting both criteria (blue; PIGF-2, PROLIFERIN, PENTRAXIN-3). Welch's t-test with Benjamini–Hochberg FDR correction; n = 2 mice per genotype.

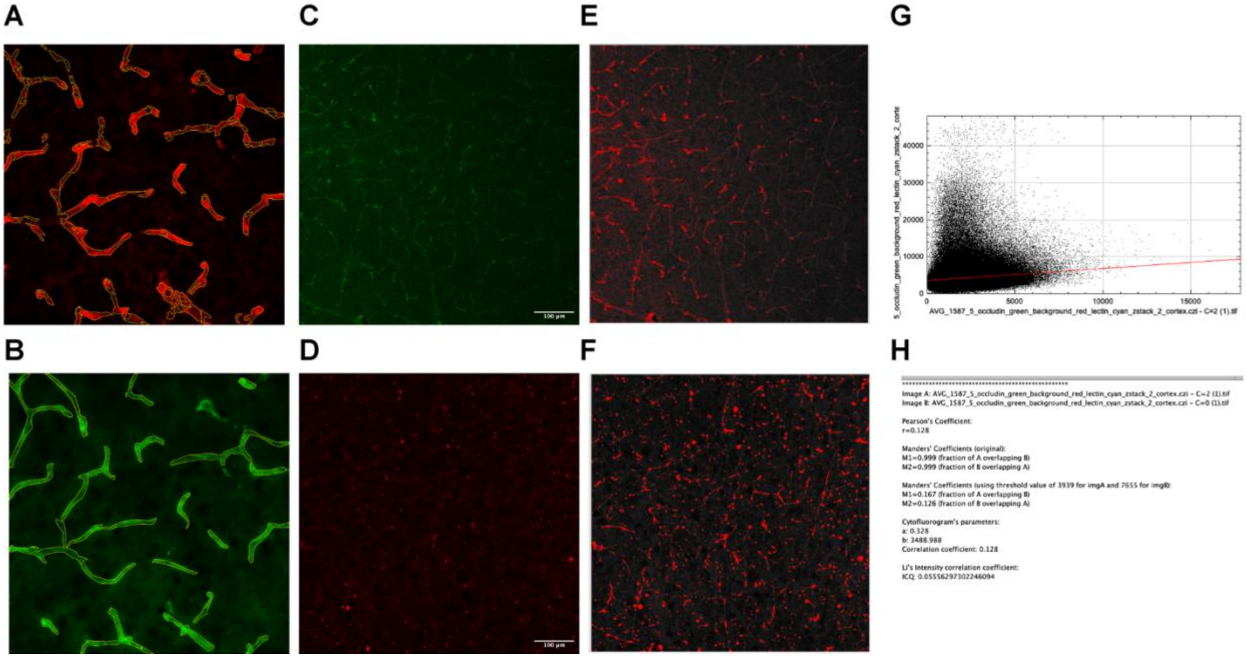


**Supplementary Figure 3. Image-analysis methods used for fluorescence colocalization. (A–B)** Manual area-based colocalization in ImageJ, exemplified using AQP4 (red) and CD31 (green) immunostaining of cortical microvessels. Vessel segments were outlined in the CD31 channel using the polygon selection tool and added to the ROI manager; the area of co-localizing AQP4 signal within each ROI was then measured. The same manual workflow was applied to caldesmon/CD31 (pericyte coverage) image pairs. **(C–H)** Semi-automated pixel-based colocalization in ImageJ using the JACoP plugin, exemplified using Occludin (red) and CD31 (green) immunostaining. **(C, D)** Background-thresholded single-channel inputs and **(E, F)** merged previews used for JACoP image preparation. **(G)** Cytofluorogram (red versus green pixel-intensity scatter plot) generated by JACoP. **(H)** JACoP numerical output reporting Pearson's correlation coefficient and Manders' overlap coefficients (M1, M2); Pearson's r values were used as the colocalization metric for the Claudin-5/lectin and Occludin/lectin analyses in Figure 4. Scale bar, 100 µm. The example image used here is the same as the one in the main Figure 6E.

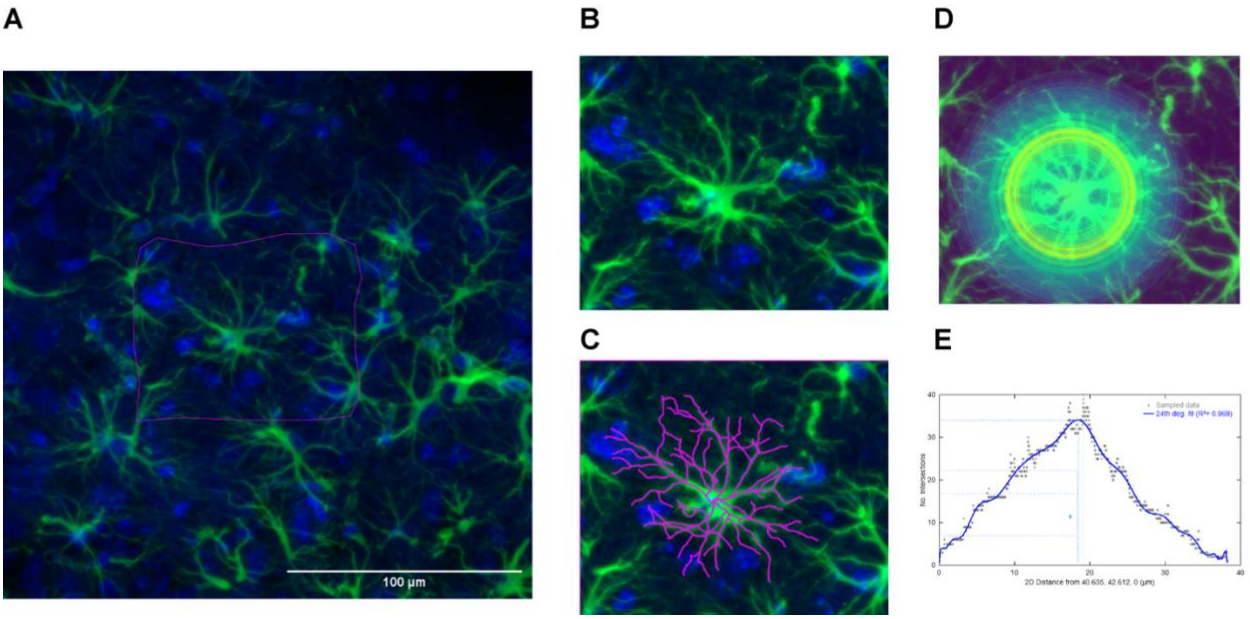


**Supplementary Figure 4: Image-analysis pipeline for astrocyte branching, length, number, and Sholl analysis. (A)** Representative cortical field of GFAP (green) and DAPI (blue) immunostaining used for individual-astrocyte selection. **(B)** Higher-magnification crop of a single astrocyte. **(C)** Simple Neurite Tracer (SNT) tracing of the same astrocyte (magenta), with the DAPI nucleus set as the soma anchor and primary and secondary branches traced outward; tracings were used to compute total branch length and number per cell. **(D)** Sholl analysis with concentric circles of stepwise-increasing radius drawn from the soma. **(E)** Representative Sholl-intersection profile (intersections versus radial distance) used to quantify branching complexity. See Methods, *Image analysis: Astrocyte Branching Pattern Analysis*. Scale bar, 100 µm. The example image used here is the same as the one in the main Figure 6A.

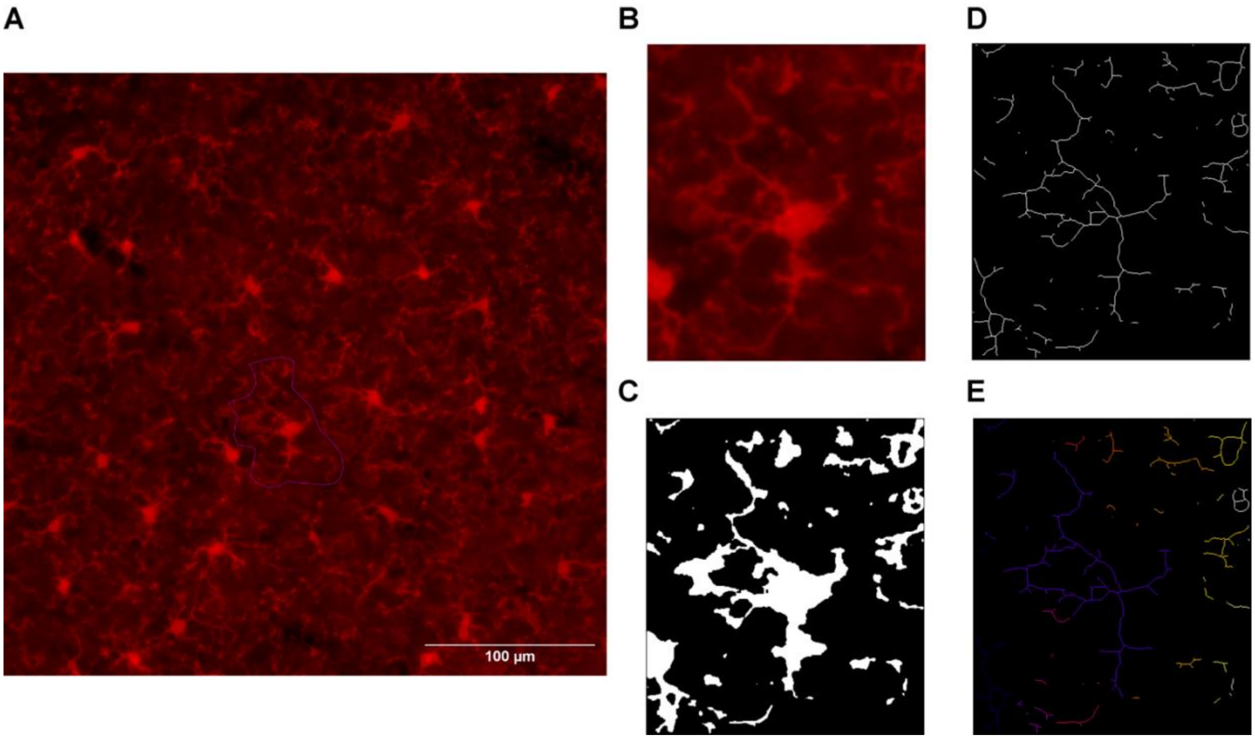


**Supplementary Figure 5. Image-analysis pipeline for microglial morphology quantification. (A)** Representative cortical Iba-1 image with the randomization grid overlay (custom ImageJ macro) used to subdivide each field into squares approximately the size of a single microglial cell; three of five randomly selected microglia were retained for analysis. **(B)** Higher-magnification crop of an individual Iba-1$^{+}$ microglial cell. **(C)** Binarized image after thresholding, brush-tool cleanup, and despeckle/denoise/remove-outliers filtering. **(D)** Skeletonized output. **(E)** Analyze Skeleton output (slab pixels, branch points, and end points color-coded), from which branch number, branch length, and junction count were quantified; cell-body area was measured separately on the original crop. See Methods, *Image analysis: Microglial Morphology Analysis*. Scale bar, 100 µm.
